# Super-Resolution Imaging Uncovers Nanoscale Tau Aggregate Hyperphosphorylation Patterns in Human Alzheimer’s Disease Brain Tissue

**DOI:** 10.1101/2024.04.24.590893

**Authors:** Adriana N. Santiago-Ruiz, Siewert Hugelier, Charles R. Bond, Edward B. Lee, Melike Lakadamyali

**Author notes:** Correspondence should be sent to M.L.

## Abstract

Tau aggregation plays a critical role in Alzheimer’s Disease (AD), where tau neurofibrillary tangles (NFTs) are a key pathological hallmark. While much attention has been given to NFTs, emerging evidence underscores nano-sized pre-NFT tau aggregates as potentially toxic entities in AD. By leveraging DNA-PAINT super-resolution microscopy, we visualized and quantified nanoscale tau aggregates (nano-aggregates) in human postmortem brain tissues from intermediate and advanced AD, and Primary Age-Related Tauopathy (PART). Nano-aggregates were predominant across cases, with AD exhibiting a higher burden compared to PART. Hyperphosphorylated tau residues (p-T231, p-T181, and p-S202/T205) were present within nano-aggregates across all AD Braak stages and PART. Moreover, nano-aggregates displayed morphological differences between PART and AD, and exhibited distinct hyperphosphorylation patterns in advanced AD. These findings suggest that changes in nano-aggregate morphology and hyperphosphorylation patterns may exacerbate tau aggregation and AD progression. The ability to detect and profile nanoscale tau aggregates in human brain tissue opens new avenues for studying the molecular underpinnings of tauopathies.

## Introduction

Tau is a microtubule-associated protein that is intrinsically disordered, highly soluble and is prominently found in neuronal axons (*1–3*). It plays a crucial role in the assembly and stability of microtubules (*1–3*) and in regulating intracellular transport by motor proteins (*4–7*). However, in a group of neurodegenerative diseases collectively known as tauopathies including Alzheimer’s Disease (AD), tau becomes mislocalized and forms large, insoluble fibrillar inclusions within the somatodendritic compartment (*1–3, 8–11*). One of the neuropathological hallmarks of AD is the formation of neurofibrillary tangles (NFTs), which are composed of aggregated tau protein (*2,12, 13*). In their seminal work, Braak and colleagues were the first to demonstrate through silver staining that NFTs spread in a stereotypic pattern across the brain (*13*). The spread of NFT-positive staining begins in the transentorhinal region of the hippocampus (Braak stage I), progresses to the subiculum region of the hippocampal pyramidal cell layer (Braak stage II), moves to the entorhinal cortex and the hippocampal pyramidal cell layer (Braak stage III), and then to the superior temporal cortex and frontal cortex (Braak stage IV) (*13–16*). At later AD stages, the staining intensifies in the hippocampus, extends to the neocortex, and eventually reaches the primary cortical areas (Braak stages V – VI) (*13–16*). Braak stages V – VI are strongly associated with clinically observable dementia, whereas Braak stages I – II typically occur in individuals who are clinically asymptomatic (*13–16*). Based on the correlation between Braak stages and clinical manifestations, it is suggested that tau aggregates play a significant role in the progression of AD (*2,12,17*).

Tau undergoes various post-translational modifications (PTMs), such as glycosylation, acetylation, and phosphorylation, which are important in regulating both its normal functions and its propensity to aggregate in disease (*10,18,19*). Notably, tau has more than 70 phosphorylation-prone residues and NFTs isolated from human AD brain tissues consist of tau that is hyperphosphorylated, often referred to as phospho-tau or p-tau (*20–22*). Hyperphosphorylation means tau has a higher degree of phosphorylation than normal, including at residues that under healthy physiological conditions are unphosphorylated or lowly phosphorylated (*22,23*).

The Proline-Rich Region (PRR) of tau, located just before the Microtubule-Binding Region (MTBR) and near the N-terminus, is particularly susceptible to hyperphosphorylation (*24*). This region contains several hyperphosphorylated residues relevant to tauopathies and that are recognized by highly specific p-tau antibodies (*25*). For example, hyperphosphorylated Serine 202 and Threonine 205 (p-S202/T205) are dually recognized by the widely used antibody AT8 (*26*). AT8 staining revealed that p-S202/T205 is primarily observed in NFTs, whereas it is less evident in the pre-NFT stage, where aggregates are non-fibrillar and appear as punctate formations (*27,28*). Given the presence of p-S202/T205 in NFTs, AT8 staining has replaced silver staining as a method for staging tau pathology in AD (*29*). While AT8 staining appears throughout all stages of AD, the highest and most significant staining is observed at the most advanced neuropathologic stage (Braak stages V – VI) (*28*). Given these observations, the dual p-S202/T205 is sometimes considered to be a marker of late-stage tau aggregation. In contrast to p-S202/T205, hyperphosphorylation at Threonine 231 (p-T231), which is recognized by AT180 antibody (*30,31*), is associated with pre-NFTs (*27*) and is significantly increased in certain brain regions at earlier Braak stages (Braak stages III – IV) (*28*). Moreover, p-T231 shows a progressive increase across Braak stages (*28*). As a result, p-T231 is sometimes referred to as a marker of early-stage tau aggregation.

The prevalence of p-tau in AD and other tauopathies has raised the possibility of using p-tau as a disease biomarker (*32*). Cross-sectional analysis of cerebral spinal fluids (CSF) and plasma samples from different clinically diagnosed AD patients have identified several potential biomarkers including p-T231, p-T217 and p-T181 (*33–35*). For example, tau phosphorylated at Threonine 181 (p-T181), also found within tau’s PRR, is increased in pre-clinical AD stage, when only subtle amyloid-β pathology exists (*32,33*). Given that tau becomes progressively hyperphosphorylated in disease and has large potential for combinatorial hyperphosphorylation, recent work further explored the mechanisms behind multi-residue tau phosphorylation. Interestingly, this work revealed several “master” residues, including p-T181, whose phosphorylation governs subsequent multi-residue tau phosphorylation (*36*). Overall, these studies highlight the disease relevance of several p-tau residues within tau’s PRR region.

For many years, NFTs and other micron-sized (> 500 nm) aggregates that are easily detectable in human brain tissues using either silver staining or p-tau antibody staining, and light microscopy were thought to be the toxic tau aggregates. However, in-vitro and cell-based experiments have demonstrated that nano-sized (< 250 nm) tau aggregates including pre-NFT tau aggregates, small fibrils and particularly soluble tau oligomers, seed the aggregation of tau into larger insoluble polymorphous fibrillar species, some of which resemble the NFTs detected in human AD brain tissues (*37–40*). Further studies in which cell and mice models were exposed to different in-vitro generated tau oligomers or tau fibrils extracted from human AD brain tissues and other tauopathies revealed that tau oligomers can recapitulate tau’s pathologic behavior associated with a specific tauopathy (*41–45*). Mouse models expressing inducible aggregation-prone human tau develop cognitive defects (*46*). Interestingly, these defects can be rescued by halting the expression of soluble tau, even when insoluble tau aggregates remain (*46*). This suggests that it is the soluble, rather than the insoluble, tau aggregates that contribute to cognitive decline. Tau oligomers were also shown to impair memory and cause synaptic dysfunction in wild type mice (*47*) and targeting tau oligomers prevented cognitive impairment and tau toxicity (*48*). Furthermore, it is thought that tau aggregation spreads in a prion-like manner through the uptake of seed-competent, oligomeric tau by neighboring cells (*49–52*). Consequently, it is currently believed that pre-NFT tau aggregates, especially tau oligomers, are the primary toxic species that contribute to the initiation of tau aggregation and subsequent neurodegeneration across different tauopathies (*53–55*).

Notably, p-T231 is enriched in tau oligomers, and p-T231 as well as p-T181 correlate with tau multimerization (*56*). Furthermore, Mass Spectrometry analyses of tau aggregates extracted from brains of AD patients have revealed variability in the levels of p-tau and oligomeric tau among patients (*57,58*). The variability in soluble and oligomeric tau as well as p-T181 correlated with the tau seeding activity and clinical outcomes (*57*). Taken together, these studies highlight the potential significance of certain PTMs, including p-T231 and p-T181, in identifying toxic oligomeric species.

Recent advancements have enabled the characterization of insoluble tau aggregates from human AD brain tissues and other tauopathies at the atomic level using Cryo-Electron Microscopy (Cryo-EM) (*59,60*). Cryo-EM analysis has revealed that distinct tau inter-protofilament packing contributes to the formation of paired helical filaments and straight filaments (*60*). Cryo-EM combined with Mass Spectrometry based proteomics revealed that tau PTMs may mediate inter-protofilament interfaces (*61*). These findings strongly suggest a significant relationship between the PTM profiles of tau aggregates, the structure of tau protofilaments, and their evolution within and across different tauopathies. However, the approaches employed so far rely on extracting tau aggregates from brain tissue, making it impossible to determine the PTM profile of tau proteins within individual tau aggregates *in-situ*. Furthermore, most studies have focused on large insoluble tau aggregates and the molecular nature of individual oligomers, including their PTM profiles, is still an open question.

Super-resolution microscopy is an advanced light microscopy method that overcomes the diffraction limit of conventional light microscopy (∼250nm), enabling the visualization and characterization of biological complexes with sizes well-below this limit (*62*). Recently, we utilized super-resolution imaging to examine tau aggregates in an engineered cell model designed to mimic tau’s aggregation seen in tauopathies (*63*). This model expresses tau harboring a P301L point mutation, a genetic mutation linked with the tauopathy Fronto-temporal Dementia with Parkinsonism linked to Chromosome 17 (FTDP-17) (*64*). Super-resolution images of the cell model under non-aggregated conditions revealed a “patchy” distribution of tau along the microtubules (*63*). Further quantitative analysis identified these patches as microtubule-bound tau monomers, dimers, and trimers (*63*). In contrast, super-resolution images of the cell model in aggregated conditions revealed the presence of cytosolic higher-order tau species such as oligomers containing four or more tau proteins, and insoluble polymorphous aggregates (small fibrils and NFT-like aggregates) (*63*). This work shows that super-resolution light microscopy can be employed to detect nano-sized tau aggregates such as tau oligomers and small fibrils in engineered cell models. However, it still remains unclear if these nanometric tau aggregates can be detected and evaluated in human brain tissue.

Here, we employed DNA-PAINT super-resolution imaging (*65*) to visualize and quantitatively analyze the extent of three disease relevant p-tau residues (p-T231, p-T181, p-S202/T205) in nano-sized (tau oligomers and small fibrils) and micron-sized (large fibrils and NFTs) tau aggregates within the temporal lobe of human postmortem brain tissues from AD patients at different Braak stages (Braak IV and Braak VI). We compared the characteristics of p-tau aggregates in AD brains to those found in a non-demented individual neuropathologically diagnosed with Primary Age-Related Tauopathy (PART-Braak stage II), a condition commonly observed in aging brains and characterized by the presence of tau pathology but lacking amyloid-beta pathology (*66*). Our findings reveal that nano-sized tau aggregates, which we refer to as nano-aggregates, are the most prevalent species of tau aggregates across AD-Braak stages and in PART, accounting for over 50% of all tau aggregates. These results highlight the sensitivity of DNA-PAINT imaging in identifying nano-sized p-tau aggregates in a brain region devoid of histological tau pathology at the earliest Braak stage, even before the onset of clinically observable symptoms and neuropathological changes associated with AD. We observed the presence of “early” (p-T231, p-T181) and “late” (p-S202/Thr205) hyperphosphorylated tau residues within tau nano-aggregates throughout all cases. Additionally, our findings reveal that the burden of tau nano-aggregates drastically increases in AD compared to PART, although it remains consistent across different AD-Braak stages. Moreover, we noted the emergence of intermediate-sized and micron-sized aggregates in AD but not in PART. The burden of these larger aggregates increased with advancing AD-Braak stages. Utilizing morphometric analysis with our newly developed algorithm, ECLiPSE (*67*), we discovered that tau nano-aggregates exhibited unique morphological characteristics between PART and AD. Additionally, multi-color DNA-PAINT imaging revealed that tau nano-aggregates display specific combinatorial hyperphosphorylation patterns at different AD-Braak stages, suggesting that variations in both morphology and hyperphosphorylation patterns of these nano-aggregates may contribute to the acceleration of tau aggregation and the advancement of the disease. Our method should be broadly applicable for characterizing the PTM profiles of tau aggregates across many tauopathies.

## Results

### Quantitative super-resolution microscopy pipeline for visualizing tau aggregates in postmortem human brain tissue

To visualize tau aggregates, we utilized formalin-fixed and paraffin embedded human postmortem brain tissues from the University of Pennsylvania (Penn) Center for Neurodegenerative Disease Research (CNDR). Specifically, we used 6 µm thick sections of the temporal cortex from three distinct cases (see **Table 1**, **Figure 1A** and **Supplementary Figure 1**). The first case, Case A, is a 74-year-old male clinically diagnosed with Parkinson’s Disease without signs of dementia. Pathologically, this case was identified as brain stem predominant Lewy Body Disease and Primary Age-Related Tauopathy (PART) with a neuropathologic change classification of A0, B1, C0, equating to Braak-stage II (**Figure 1A**). PART is commonly observed in aging brains and is characterized by the presence of tau aggregates and absence of amyloid-beta pathology, therefore not consistent with the neuropathologic designation of Alzheimer’s Disease (ADNC). Case B is a 97-year-old male clinically normal without signs of dementia, but pathologically showing an intermediate level of ADNC (A2, B2, C1), which we refer to as intermediate AD or AD-Braak stage IV (**Figure 1A**). The third case, Case C, is a 68-year-old male clinically diagnosed with dementia and probable AD, and pathologically presenting a high level of ADNC (A3, B3, C3), which we refer to as advanced AD or AD-Braak stage VI (**Figure 1A**).

**Figure 1:**
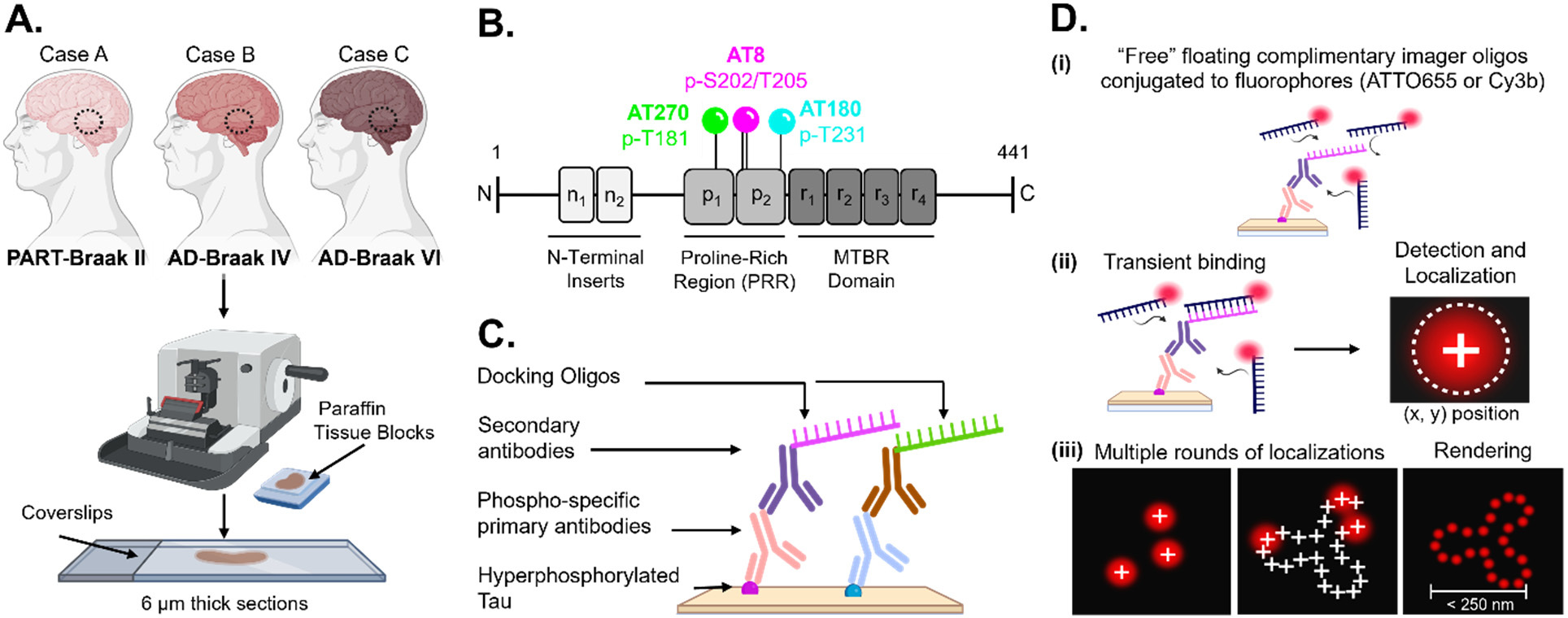
Schematic representation of the preparation and imaging of postmortem human brain tissue sections. **(A)** Temporal cortex tissue blocks from three cases neuropathologically diagnosed as Primary Age-Related Tauopathy (PART-Braak II), Alzheimer’s Disease Braak stages IV and VI (AD-Braak IV and VI, respectively), were sectioned to a 6 μm thickness and mounted onto treated coverslips. **(B)** Sections were immunolabeled with validated phospho-specific primary antibodies AT180, AT8, and AT270 to target disease-confirmed hyperphosphorylated tau residues Threonine 231 (p-T231), Serine 202/Threonine 205 (p-S202/T205), and Threonine 181 (p-T181), respectively. All sites are located in Tau’s Proline-Rich Region (PRR). **(C)** After incubation with primary antibody, a secondary antibody conjugated to a unique DNA-sequence (docking oligo) is added. **(D)** Super-resolution images are acquired by performing DNA-PAINT. The schematic diagram shows a summary of the imaging process: (i) immunolabeled samples are incubated with a solution of an imager oligo (imager-probes), which is a fluorophore-conjugated (ATTO655 or Cy3b) DNA-sequence complementary to the docking oligos; (ii) these “free” floating imager oligos will transiently bind to the docking oligo, allowing for the detection of a signal; (iii) this information is then utilized to calculate the relative position of that fluorophore in space (x, y position) through a process known as localization. The cycle is then repeated for a number of frames to accumulate a collection of localizations. At the end of the acquisition, all these frames are merged to render a high-resolution image of the labeled object of interest, such as tau aggregates.

**Table 1:** Information on the postmortem human brain tissues used in this study.

| Case | Sex | Age | Clinical Diagnosis | Pathological Diagnosis | ADNC Score | Braak Stage |
| --- | --- | --- | --- | --- | --- | --- |
| A | Male | 74 | Parkinson's Disease | Brainstem-Predominant<br>Lewy Body Disease<br>-----<br>Primary Age Related<br>Tauopathy | A = 0; B = 1; C = 0 | II |
| B | Male | 97 | Normal | Intermediate level of<br>Alzheimer's Disease | A = 2; B = 2; C = 1 | IV |
| C | Male | 68 | Probable Alzheimer's<br>Disease | High level of<br>Alzheimer's Disease | A = 3; B = 3; C = 3 | VI |

We chose the temporal cortex for our studies because NFT pathology, a prominent hallmark of AD, is detectable in this region during the disease’s intermediate stages (Braak stages III – IV). The tissue samples were sectioned to a 6 µm thickness as we determined it to be the optimal thickness for visualizing tau aggregates with minimal background (see Methods). Prior to immunostaining, samples were treated with Xylene and Ethanol to deparaffinize and rehydrate the tissues. Afterwards, we performed a heat-induced antigen retrieval, followed by immunostaining with phospho-specific antibodies (see Methods). To reduce background and optical aberrations, we included a clearing step with 2,2-thiodiethanol (see Methods).

For immunostaining, we used commercially available and previously validated (*68*) primary phospho-specific antibodies: AT180, AT8, and AT270, targeting hyperphosphorylated tau residues p-T231, p-S202/T205, and p-T181, respectively (**Figure 1B and Tables 2 and 3 in Methods**). These phospho-specific antibodies were labeled with secondary antibodies conjugated to unique orthogonal DNA-docking oligos, facilitating DNA-PAINT super-resolution microscopy (see Methods and **Figure 1C and D**).

**Table 2.** List of primary antibodies.

| Antibody | Target | Host Species | Catalog Number | Vender |
| --- | --- | --- | --- | --- |
| PThr231 | p-Thr231 | Rabbit (monoclonal) | SAB4504563 | Sigma-Aldrich |
| AT180 | p-Thr231 | Mouse (monoclonal) | MN1040 | Invitrogen |
| AT270 | p-Thr181 | Mouse (monoclonal) | MN1050 | Invitrogen |
| D9F4G | p-Thr181 | Rabbit (monoclonal) | 12885 | Cell Signaling |
| AT8 | p-Ser202,Thr205 | Mouse (monoclonal) | MN1020 | Fisher Scientific |

**Table 3.** List of secondary antibodies and imaging probes.

| Antibody and Imager Probes | Target | Catalog Number | Vendor |
| --- | --- | --- | --- |
| Anti-mouse IgG + Docking Site 1 | Mouse-primary antibodies | N/A | Massive Photonics |
| Anti-rabbit IgG + Docking Site 2 | Rabbit-primary antibodies | N/A | Massive Photonics |
| Imager-probe 1: ATTO 655 | Mouse IgG docking site 1 | N/A | Massive Photonics |
| Imager-probe 2: Cy3b | Rabbit IgG docking site 2 | N/A | Massive Photonics |
| Donkey anti-mouse Alexa-647 | Mouse-primary antibodies | A31571 | Invitrogen |
| Donkey anti-rabbit Alexa-546 | Rabbit-primary antibodies | A10040 | Invitrogen |

Super-resolution DNA-PAINT imaging revealed hyperphosphorylated tau proteins within tau aggregates in both PART and AD across all three examined residues (**Figure 2A-E**). In each case (p-T231, p-S202/T205, and p-T181), we observed an increase in the abundance, size, and morphological complexity of tau aggregates corresponding with higher Braak stages (**Figure 2B-D**). Specifically, tau aggregates in PART-Braak stage II were nanometric and punctate, resembling the tau oligomers we previously observed in engineered cell models using super-resolution microscopy (*63*). In contrast, AD brain samples not only contained these nanometric and punctate aggregates, but we also observed larger aggregates, including linear fibrillar structures and amorphous aggregates resembling NFTs (**Figure 2B-D**). To confirm that the structures we observed corresponded to tau aggregates rather than background and noise, we carried out negative control experiments using brain tissues from PART-Braak stage II and AD-Braak stage VI, immunolabeled only with secondary DNA-PAINT antibodies (without primary phospho-tau antibodies) and imaged under identical conditions as the positive samples (**Figure 2A,E**). We specifically chose PART (**Figure 2A**) and advanced AD (**Figure 2E**) brain tissues for these negative control experiments as these conditions correspond to the lowest and highest tau aggregate burden, respectively. These negative controls enabled us to assess the background signal arising from non-specific secondary antibody labeling and off-target interactions from imager oligos in DNA-PAINT images. The negative controls exhibited significantly fewer detected localizations compared to positive samples, and the background puncta were also smaller in size than the tau aggregates visualized in positive samples (compare **Figure 2A,E** to **Figure 2B-D**).

**Figure 2:**
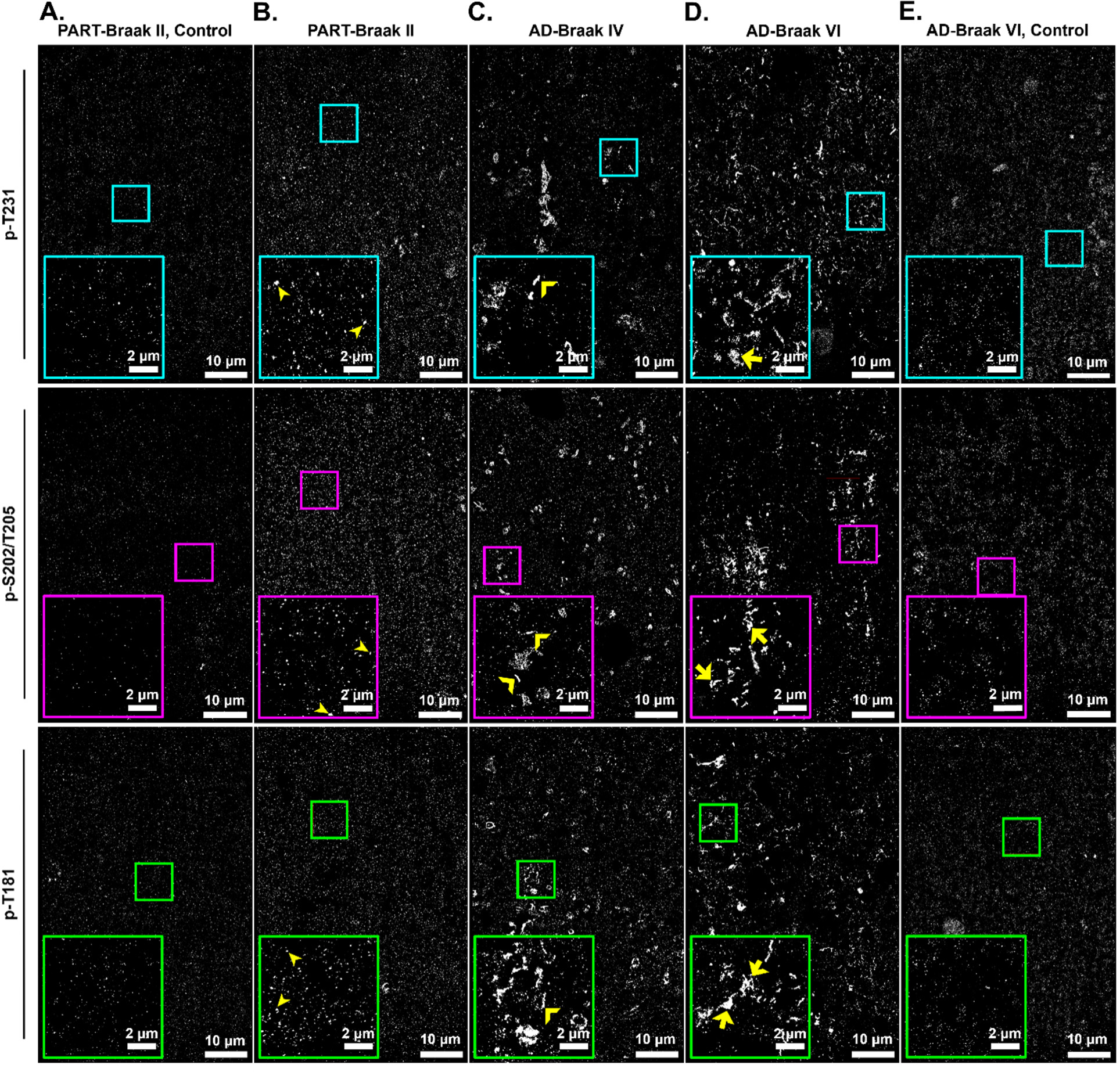
Single-color DNA-PAINT images of tau aggregates in postmortem human brain tissues. **(A,E)** Representative single-color DNA-PAINT images of negative control samples. Sections from PART-Braak II **(A)** or AD-Braak VI **(E)** were immunolabeled and imaged using the pipeline described in Figure 1, but in the absence of phospho-specific primary antibodies. Inset shows sparse background localizations. **(B)** Representative single-color DNA-PAINT images of hyperphosphorylated tau in tissue sections from PART-Braak II immunolabeled with AT180, AT8, and AT270 phospho-specific antibodies. Insets reveal nano-sized, punctate tau aggregates (filled arrowheads) that are larger in size than those present in negative control images. **(C)** Representative single-color DNA-PAINT images of hyperphosphorylated tau in tissue sections from AD-Braak IV immunolabeled with AT180, AT8, and AT270 phospho-specific antibodies. Inset shows the presence of an array of tau aggregates such as linear fibrils (empty arrowhead). **(D)** Representative single-color DNA-PAINT images of hyperphosphorylated tau in tissue sections from AD-Braak VI immunolabeled with AT180, AT8, and AT270 phospho-specific antibodies. Insets reveal the presence of an array of tau aggregates such as amorphous aggregates (arrows). **(A-E)** AT180 targets (p-T231) were imaged with imager-probe 2 (Cy3b). AT8 targets (p-S202/T205) were imaged with imager-probe 1 (ATTO655). AT270 targets (p-T181) were imaged with imager-probe 2 (Cy3b).

To quantify the abundance, size, and morphology of phospho-tau aggregates in DNA-PAINT super-resolution images, we developed a quantitative analysis pipeline (**Figure 3**). We applied a commonly utilized segmentation strategy for super-resolution microscopy, Voronoi Tessellation (*63,69*), to segment individual tau aggregates in all positive samples as well as the background/noise present in DNA-PAINT images of negative control samples (see Methods and **Figure 3A-C**). To remove the background and noise from the images of positive samples (**Figure 3D,E**), we plotted and compared the area distribution of the segmented objects in the images of positive samples and negative controls (**Figure 3F**, Supplementary Figure 2A,B). We found that tau aggregates in the positive samples had a median area significantly larger than the background/noise in the negative controls (**Figure 3F, Supplementary Figure 2A,B**). To remove background/noise from the positive samples, we applied an area threshold (indicated by the red dashed line in **Figure 3F and Supplementary Figure 2A,B**), effectively excluding the majority (>97%) of background/noise structures corresponding to those segmented in the negative controls (**Figure 3D,E**).

**Figure 3:**
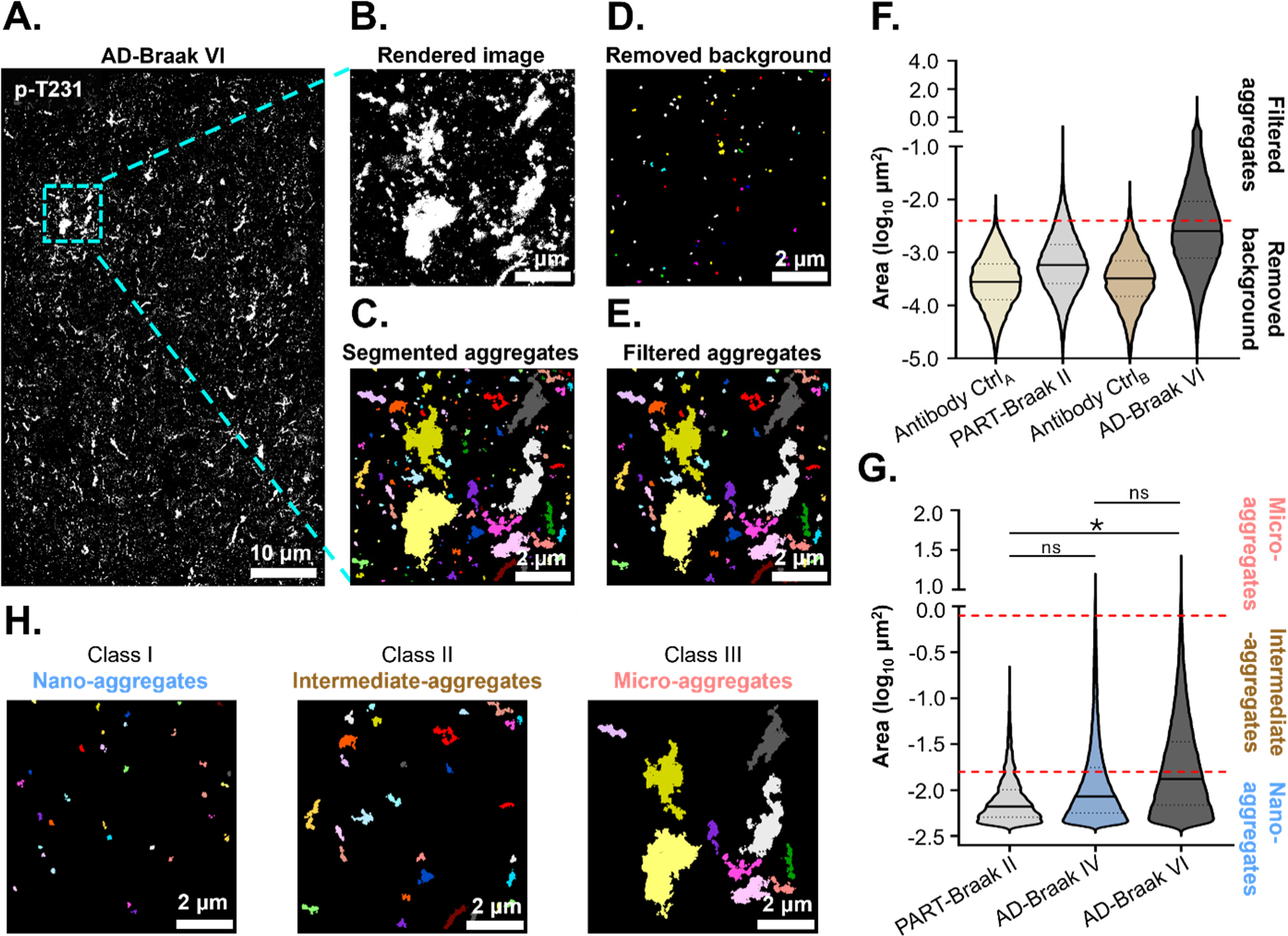
Quantitative analysis pipeline of tau aggregates. **(A)** Representative single-color DNA-PAINT image of p-T231 in AD-Braak VI section, immunolabeled with AT180 antibody and imaged with imager-probe 2 (Cy3b). **(B)** Inset shows the image rendered from point localizations of p-T231 tau (white). **(C)** The point localizations are clustered and segmented using Voronoi Tessellation. Inset image shows corresponding Voronoi segmentation. Segmented image is pseudo-color coded with different colors corresponding to different tau aggregates. **(D-E)** Insets show the removed background signal **(D)** and the filtered tau aggregates after background removal **(E)**. **(F)** Violin plot showing the area distribution of segmented objects from the negative control images (Antibody Ctrl_A_ = PART-Braak II, Antibody Ctrl_B_ = AD-Braak VI) and positive control samples (PART-Braak II and AD-Braak VI). Red dashed line indicates the area cut-off (−2.4 log_10_ μm^2^= 0.004 μm^2^) used for the area-based filtering of background signal. Segmented objects below this value are removed from the list of segmented tau aggregates (insets D and E). Solid line indicates median, and the dotted lines indicate the 25th and 75th percentile. Antibody negative controls (A: Part-Braak II; B: AD-Braak VI) N = 1 tissue section, n = 4 fields of view; PART-Braak II: N = 4 tissue sections, n = 15 fields of view; AD-Braak VI: N = 4 brain tissue slices, n = 17 fields of view. **(G)** Violin plots showing the area (log10 μm^2^) per tau aggregates after background filtering for p-Thr231 images (light grey: PART-Braak II sections; blue grey: AD-Braak IV tissue sections; dark grey: AD-Braak VI tissue sections). All sections were immunolabeled with AT180 antibody and imaged with imager-probe 2 (Cy3b). Red dashed lines indicate the area-cut off used for classifying tau aggregates: nano-aggregates (0.004 – 0.017 μm^2^), intermediate-aggregates (0.017 – 0.15 μm^2^) and micro-aggregates (above 0.15 μm^2^). Solid line indicates median, and the dotted lines indicate the 25th and 75th percentile. PART-Braak II: N = 4 tissue sections, n = 15 fields of view; AD-Braak IV: N = 4 tissue sections, n = 18 fields of view; AD-Braak VI: N = 4 brain tissue slices, n = 17 fields of view. An unpaired, non-parametric Mann-Whitney test was performed between PART and AD-stages IV and VI. A p value of < 0.05 was taken as statically significant. P values = (ns) >0.05, (*) 0.05 – 0.03, (**) 0.002 – 0.03, (***) 0.0002 – 0.002, (****) 0.0001 – 0.0002. **(H)** Representative examples of tau aggregates belonging to Class I (nano-aggregates), Class II (intermediate-aggregates) and Class III (micro-aggregates).

After applying this filter, we reassessed the area distribution of tau aggregates in the filtered images from PART and AD-Braak Stages IV and VI (**Figure 3G, Supplementary Figure 2C,D**). We observed that tau aggregates were smallest in PART and increased in size, particularly in advanced AD-Braak stages (**Figure 3G, Supplementary Figure 2C,D**), reflecting the progressive aggregation process where tau oligomers seed the formation of fibrils and other larger amorphous aggregates, including NFTs. Utilizing these area distributions, we classified tau aggregates into three classes: nano-aggregates (the smallest aggregates, predominant in PART-Braak stage II), intermediate-aggregates (mid-sized aggregates, primarily found in AD-Braak stages IV and VI), and micro-aggregates (the largest aggregates, appearing in the tail-end of the area distribution for AD-Braak stages IV and VI) (**Figures 3G,H, Supplementary Figure 2C,D**). Further visual inspection of tau aggregates within these classes confirmed a progression in size and morphology, starting with small, punctate nano-aggregates (class I) and evolving to larger, more complex aggregates (classes II and III) (**Figure 3H**).

Overall, we successfully established a quantitative super-resolution microscopy pipeline that effectively identifies phospho-tau-enriched tau aggregates, ranging from nano- to micron-sized, in PART and AD postmortem human brain tissue.

### Nano- to micro-aggregate burden increases in AD compared to PART with unique nano-aggregate morphology emerging in AD

To determine whether the abundance of nano-, intermediate-, and micro-aggregates varies across AD stages and if this is correlated to the hyperphosphorylation of specific residues, we quantified the number of tau aggregates per unit area in the brain tissues imaged (**Figure 4A-C**). As expected, almost no tau aggregates of any size-based class were found in the negative control samples (**Figure 4A-C**). A notable increase in the number of nano-aggregates, particularly those with p-T231 (mean fold change of 36 compared to negative control) and p-T181 (mean fold change of 56 compared to negative control), was observed for PART when compared to the negative controls (**Figure 4A**). However, intermediate- and micro-aggregates were completely absent in PART (**Figures 4B,C**), aligning with the expectation that NFTs are not detectable in the temporal cortex until intermediate-Braak stages (Braak stage III onward). Interestingly, our findings reveal that by using DNA-PAINT, it is possible to detect nano-aggregates enriched with phospho-tau in brain regions devoid of significant tau pathology, which may represent physiological tau oligomers or the earliest stages of tau aggregation.

**Figure 4:**
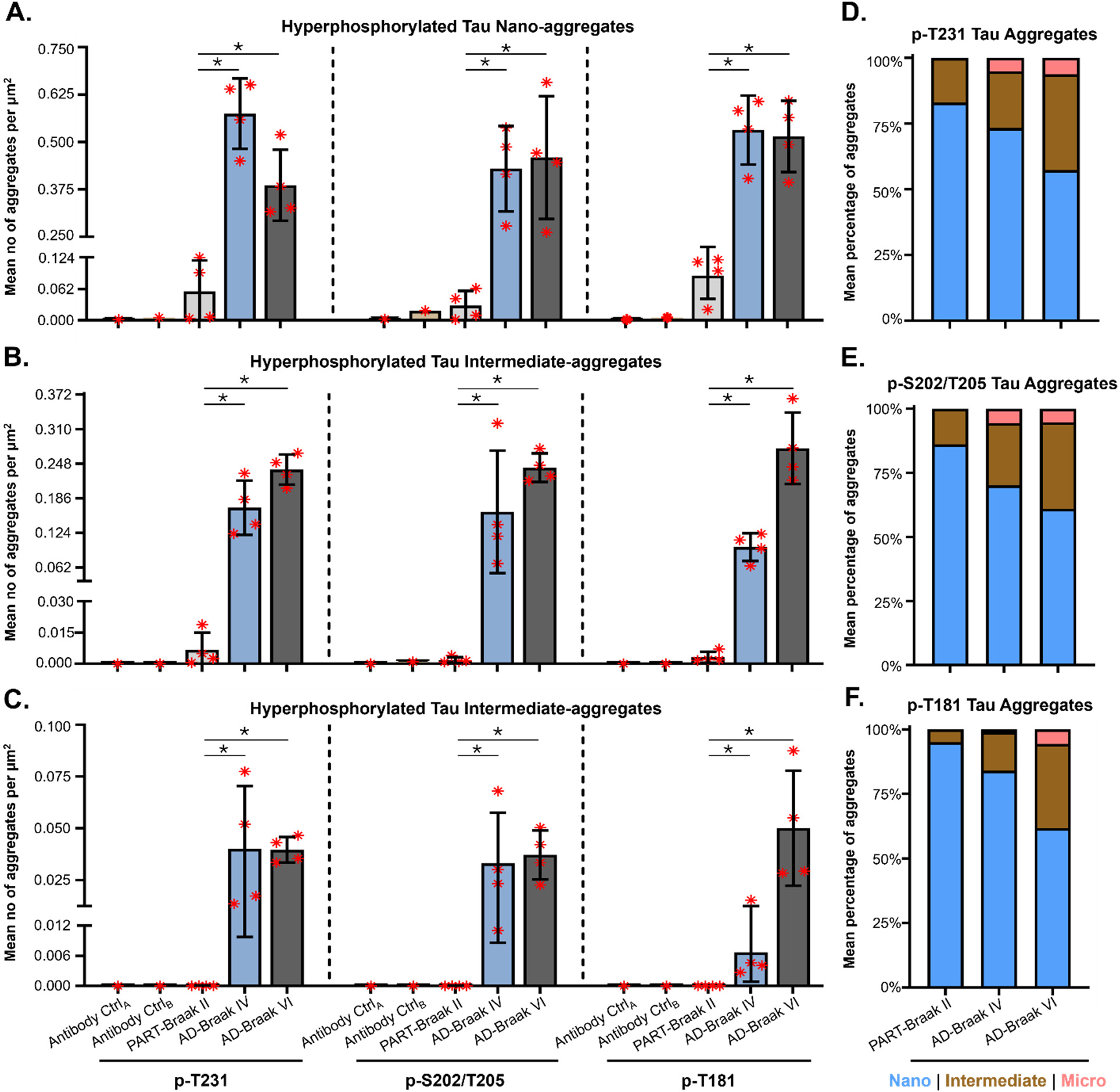
Nano-aggregates are the predominant p-tau aggregate species in PART and AD. **(A-C)** Quantification of the number of tau nano- **(A)** intermediate- **(B)** and micro-aggregates **(C)** per unit area in antibody negative control samples (Antibody Ctrl_A_: antibody negative control in PART-Braak II tissue and Antibody Ctrl_B_: antibody negative control in AD-Braak VI tissue) and in positive samples (light grey = PART-Braak II, blue grey = AD-Braak IV, dark grey = AD-Braak VI) for p-T231 immunolabeled with primary antibody AT180 and imaged with imager-probe 2 (Cy3b) (left), p-S202/T205 immunolabeled with primary antibody AT8 and imaged with imager-probe 1 (ATTO655) (middle), and p-T181 immunolabeled with AT270 and imaged with imager-probe 2 (Cy3b) (right). Bar plots show the mean number of tau aggregates per unit area, error bars are the standard deviation of the mean, and the red asterisks are the mean number of aggregates per unit area in individual tissue sections imaged. **p-T231:** control A: N = 1 tissue section, n = 4 fields of view; control B: N = tissue section, n = 4 fields of view; PART-Braak II: N = 4 tissue sections, n = 15 fields of view; AD-Braak IV: N = 4 tissue sections, n = 18 fields of view; AD-Braak VI: N = 4 tissue sections, n = 17 fields of view. **p-S202/T205:** control A: N = 1 brain tissue section, n = 4 fields of view; control B: N = 1 tissue section, n = 4 fields of view; PART-Braak II: N = 4 tissue sections, n = 17 fields of view; AD-Braak IV: N = 4 tissue sections, n = 17 fields of view; AD-Braak VI: N = 4 tissue sections, n = 18 fields of view. **p-T181:** control A: N = 1 brain tissue section, n = 4 fields of view; control B: N = 1 tissue section, n = 4 fields of view; PART-Braak II: N = 4 tissue sections, n = 17 fields of view; AD-Braak IV: N = 4 tissue sections, n = 19 fields of view; AD-Braak VI: N = 4 tissue sections, n = 18 fields of view. An unpaired, non-parametric Mann-Whitney test was performed between PART and AD-Braak stages IV and VI. A p value of < 0.05 was taken as statically significant. P values = (ns) >0.05, (*) 0.05 – 0.03, (**) 0.002 – 0.03, (***) 0.0002 – 0.002, (****) 0.0001 – 0.0002. **(D-F)** Percentage of p-T231 **(D)**, p-S202/T205 **(E)**, and p-T181 **(F)** positive tau aggregates per class calculated from data presented in A-C. Stacked bar plots show the mean percentage of all brain tissue slices imaged (blue = nano-aggregates; brown = intermediate-aggregates; pink = micro-aggregates).

The abundance of nano-aggregates significantly increased in AD-Braak stages IV and VI compared to PART (between 5 to 14-fold increase depending on the p-tau mark), and this pattern was consistent across all examined phospho-tau residues (**Figure 4A**). Although the nano-aggregate load did not change significantly between these two AD-Braak stages (**Figure 4A**), a marked increase in the burden of p-T181 modified intermediate- (3-fold increase) and micro-aggregates (7-fold increase) was observed for AD-Braak stage VI compared to Braak stage IV (**Figure 4B,C**), suggesting that appearance of large aggregates containing p-T181 tau may be a hallmark of more advanced AD.

We further determined the percentage of the different classes (nano-, intermediate- and micro-) of tau aggregates in PART and AD (**Figure 4D-F**). We found that nano-aggregates constitute the majority of aggregates in both PART and AD-Braak stage IV (83% and 73%, respectively). Notably, even for advanced AD (Braak stage VI), nano-aggregates account for over 50% of all phospho-tau aggregates. This pattern persisted across all visualized phospho-tau residues, suggesting that nano-aggregates contain tau hyperphosphorylated at both “early” (p-T231, p-T181) and “late” (p-S202/T205) phospho-tau residues. Although the proportion of intermediate- and micro-aggregates gradually increases through Braak stages, our data highlights that the large, micron-sized aggregates detectable by conventional immunofluorescence or histochemistry represent only a minor fraction of the total phospho-tau aggregate burden. Most aggregates remain smaller than the detection and resolution limit of these imaging techniques, underscoring the predominance of nano-aggregates in AD pathology.

Given the prevalence of nano-aggregates in PART and AD, we focused on characterizing this class of tau aggregates further to determine if there are any morphological variations across Braak stages. To this end, we utilized a new algorithm that we recently developed called ECLiPSE (Enhanced Classification of Localized Point-clouds by Shape Extraction) (*67*). ECLiPSE is an automated machine learning analysis pipeline that extracts morphological parameters and classifies structures captured through super-resolution microscopy. With ECLiPSE, we were able to determine 67 shape descriptors including geometric, boundary, skeleton, and other properties of each phospho-tau nano-aggregate. Through Principal Component Analysis (PCA), we investigated whether the morphology of nano-aggregates in PART and AD (**Figure 5A**) showed distinct clustering patterns in PCA-space. Our findings revealed that PART nano-aggregates consistently formed a separate cluster from those in the AD-Braak stages IV and VI, which showed a closer overlap in PCA-space (**Figure 5B, Supplementary Figure 3A**). This trend was true irrespective of the phospho-tau residue analyzed, but particularly pronounced for p-S202/T205 and p-T181 (**Figure 5B and Supplementary Figure 3A, respectively**). Further analysis of specific morphological features (area, circularity, and major axis) showed that nano-aggregates in AD brains were less circular, with a longer major axis and a larger area compared to those in PART (**Figure 5A, C and Supplementary Figure 3B**). These morphological changes may reflect a progression of tau pathology across different Braak stages. They might also highlight unique characteristics of nano-aggregates in PART, where they may represent more physiological oligomers compared to the pathological ones associated with AD.

**Figure 5:**
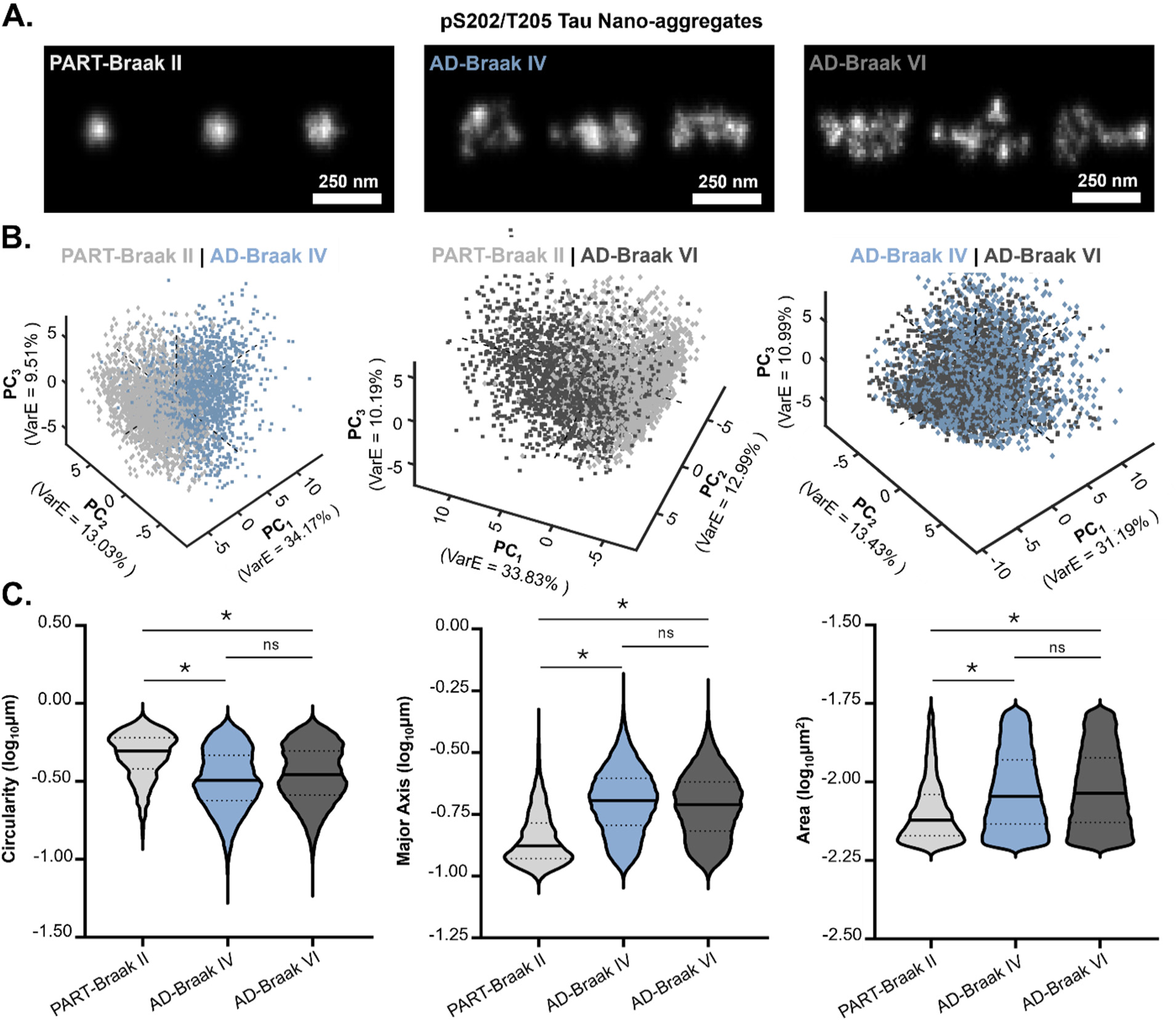
Morphological analysis of tau nano-aggregates reveals distinct characteristics between PART and AD. **(A)** Representative examples of p-S202/T205 immunolabeled nano-aggregates in PART-Braak II, AD-Braak IV and AD-Braak VI. **(B)** Principal Component Analysis (PCA) plots showing the PC 1 and 2 for p-S202/T205 immunolabeled nano-aggregates in PART-Braak II (light grey dots), AD-Braak IV (blue grey dots) and AD-Braak VI (dark grey dots). **(C)** Violin plots showing the circularity, major axis, and area for p-S202/T205 immunolabeled nano-aggregates in PART (light grey), AD-Braak IV (blue grey) and AD-Braak VI (dark grey). Solid line indicates median, and the dotted lines indicate the 25^th^ and 75^th^ percentile PART-Braak II: N = 4 tissue sections, n = 17 fields of view; AD-Braak IV: N = 4 tissue sections, n = 17 fields of view; AD-Braak VI: N = 4 tissue sections, n = 18 fields of view. A Wilcoxon rank sum test was performed between populations of tau aggregates in PART and AD-stages IV and VI. A p value of < 0.05 was taken as statically significant. P values = (ns) >0.05, (*) 0.05 – 0.01, (**) < 0.01.

Overall, our findings reveal that amongst the different species of tau aggregates, nano-aggregates are the predominant ones in both PART and AD. Furthermore, while these nano-aggregates display distinct morphological characteristics when comparing PART and AD, they remain morphologically consistent across the different AD-Braak stages (IV and VI).

### Quantitative multi-color super-resolution microscopy pipeline for visualizing the degree of multi-hyperphosphorylated residues within individual tau aggregates

Our findings revealed that tau nano-, intermediate-, and micro-aggregates consist of tau hyperphosphorylated at different residues. Tau is thought to undergo combinatorial hyperphosphorylation in disease states, with modifications at certain residues such as p-T181, facilitating further hyperphosphorylation of other tau residues (*36*). Therefore, we sought to determine whether these phospho-tau residues coexisted within the same tau aggregates in a combinatorial fashion or if they delineated separate tau aggregates. To address this question, we carried out dual-color DNA-PAINT super-resolution microscopy employing combinations of p-T181 and p-S202/T205 or p-T231 and p-S202/T205 (**Figure 6A,B**).

**Figure 6:**
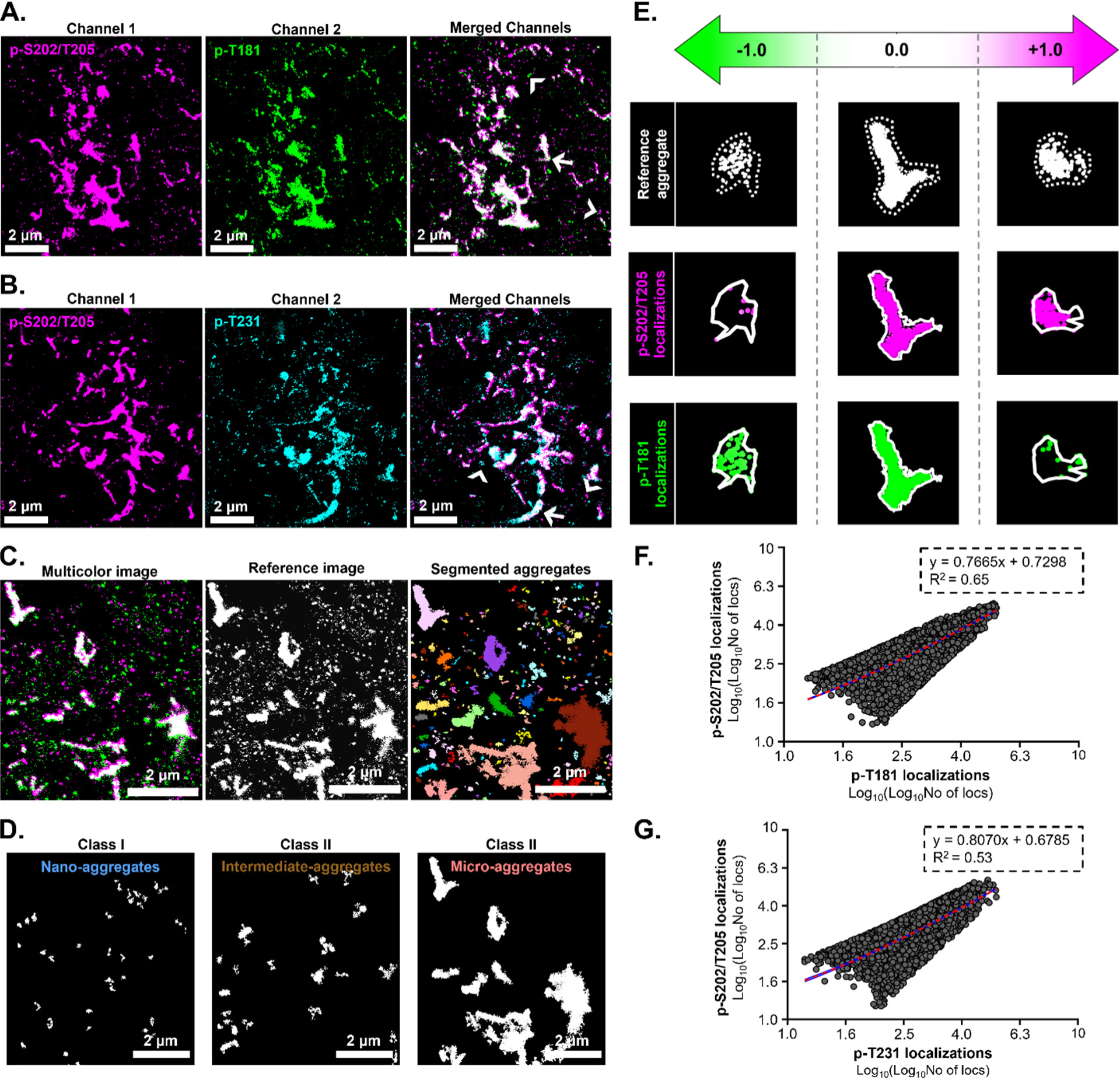
Multicolor DNA-PAINT imaging enables the identification of multiple hyperphosphorylated tau residues within a single tau aggregate. **(A)** Representative dual-color DNA-PAINT images of AD-Braak VI sections immunolabeled with primary antibodies AT8 and AT270 to identify p-S202/T205 tau (magenta) and p-T181 (green) within aggregates. Overlay reveals the presence of both targets within individual aggregates (white, arrow) as well as aggregates predominantly modified at a single target (arrowheads). **(B)** Representative dual-color DNA-PAINT image of AD-Braak VI sections immunolabeled with primary antibodies AT8 and AT180 to identify p-S202/T205 tau (magenta) and p-T231 (cyan) within aggregates. Overlay reveals the presence of both targets within individual aggregates (white, arrow) as well as aggregates predominantly modified at a single target (arrowheads). **(C-E)** Dual-color DNA-PAINT image analysis pipeline. The two channels are merged, tau aggregates are segmented **(C)**. After background filtering and quality control steps, segmented tau aggregates are split into three classes (nano-, intermediate- and macro-aggregates) **(D)**. The number of localizations belonging to each channel within the individual reference tau aggregates is computed and converted to an enrichment score, ranging from −1 to 1 **(E)**. **(F, G)** X,Y scatter plot showing the number of localizations of p-S202/T205 and p-T181 **(F)** or p-S202/T205 and p-T231 **(G)** per tau aggregate in AD-Braak VI tissue sections. Red line is the linear regression, blue shadow is the standard error. AT8 + AT270 immunolabeled AD-Braak VI tissue sections: N = 4 tissue sections, n = 16 fields of view. AT8 + AT180 immunolabeled AD-Braak VI tissue sections: N = 4 tissue sections, n = 16 fields of view.

To quantitatively analyze dual-color DNA-PAINT images, we developed a co-localization analysis pipeline (**Figure 6C-E**). First, we combined the two channels corresponding to the two modifications into a single reference channel and performed segmentation (**Figure 6C**). This ensured the segmentation of a unified set of tau aggregates from both channels, using consistent segmentation parameters. Occasionally, when two distinct tau aggregates appeared in separate channels but were spatially proximate, they might overlap partially in the merged reference image and thus be segmented as a single tau aggregate. To rectify such segmentation errors, we assessed the percentage of overlap of each individual channel within the segmented reference tau aggregate (**Supplementary Figure 4 and Methods**). A high percentage overlap suggests that the detected localizations from each channel intermingle and coincide within the segmented reference aggregate, likely indicating the presence of a single aggregate with a mixed modification profile (**Supplementary Figure 4A**). Conversely, a low percentage overlap indicates that the detected localizations from each channel remain separate within the segmented reference aggregate, suggesting two distinct aggregates that contain a single modification and that partially overlap in space (**Supplementary Figure 4B**). Therefore, we conducted this quality control step to ascertain the overlap score of segmented tau aggregates in dual-color DNA-PAINT images from PART and AD. If the overlap score fell below 30%, segmented reference aggregate is separated and the localizations from each channel are re-segmented. Following this quality control step, tau aggregates exhibited an average high overlap score of 67%, confirming the effectiveness of the segmentation.

Following segmentation and filtering, we once again classified tau aggregates into three classes: nano-, intermediate-, and micro-aggregates, utilizing the area cut-offs previously established (**Figure 6D**). In the final step, we divided the detected localizations corresponding to each reference tau aggregate back into their original two-color channels, and then determined the number of detected localizations for each phospho-tau residue (**Figure 6E**). In DNA-PAINT, the number of detected localizations scales linearly with the number of visualized target protein (*70*). Consequently, we calculated the ratio between the difference of the detected localizations for each residue and the sum of the detected localizations for each residue (see Methods) to derive a relative enrichment score for each modification. A score of −1 or +1 corresponds to tau aggregates containing only one (e.g., p-T181) or the other (e.g., p-T202/S205) phospho-tau modification, while a score of 0 designates tau aggregates with an equal number of detected localizations from each phospho-tau modification (**Figure 6E**).

It is possible that steric hindrance effects could preclude the simultaneous binding of two antibodies to adjacent modified residues. To confirm this wasn’t occurring, we assessed the number of detected localizations within tau aggregates for each modification in dual-color images from all samples (PART and AD). If steric hindrance posed a problem, we would anticipate a negative correlation between the number of detected localizations (i.e., high detection for one modification correlating with low detection for the other). However, in most cases we observed a positive correlation between the localizations detected for combinations of phospho-tau modifications (**Figure 6F,G and Supplementary Figure 5A,B**). To further validate that steric hindrance wasn’t an issue, we compared the number of detected localizations within tau aggregates in single-color and dual-color DNA-PAINT images. If steric hindrance affected antibody binding, we would expect to see a decrease in the number of detected localizations in the dual-color images compared to the single-color images. However, we did not observe such a decrease (**Supplementary Figure 5C-E**).

Taken together, these findings strongly suggest that the antibodies can bind to their respective targets simultaneously without hindering each other sterically, and dual-color imaging is robust for determining the relative amounts of hyperphosphorylated tau within individual tau aggregates.

### Nano-aggregates exhibit stage specific hyperphosphorylation patterns in Alzheimer’s Disease and combinatorically hyperphosphorylated nano-aggregates are morphologically distinct

The two-color analysis pipeline enabled us to determine the extent of phosphorylation marks corresponding to distinct tau residues within tau’s PRR domain within each individual tau aggregate (**Figure 7A**), which is not possible using bulk biochemical analysis. Nano-aggregates in intermediate AD (Braak stage IV) displayed a broad distribution of enrichment scores for both p-T181 + p-S202/T205 and p-T231 + p-S202/T205 combinations, indicating varying and heterogeneous levels of tau phosphorylation at different residues within these nano-aggregates. However, in advanced AD (Braak stage VI), we observed a peak centered at an enrichment score of zero for both combinations of phospho-tau modifications (**Figure 7A**). These findings suggest that the initial heterogeneity in the enrichment scores diminishes, with nano-aggregates containing similar amounts of both phospho-tau modifications at advanced AD stages. To further analyze the phosphorylation pattern of individual tau nano-aggregates in a simpler manner, we applied cut-offs to the enrichment scores (**Figure 7A**). Scores below −0.8 and above 0.8 were categorized as “singly modified” tau aggregates, while scores between −0.8 and 0.8 were labeled as “dually modified” tau aggregates (**Figure 7B**). This analysis revealed that the majority of tau nano-aggregates in PART and intermediate AD (Braak stage IV) were modified with only one of the phospho-tau residues visualized (**Figure 7B**). However, in advanced AD (Braak stage VI), there was a substantial increase in the percentage of tau nano-aggregates that are dually modified with both phospho-tau residues, from 11% in AD-Braak Stage IV to 54% in AD-Braak Stage VI for p-T181 + p-S202/T205 and from 8% in AD-Braak Stage IV to 35% in AD-Braak Stage VI for p-T231 + p-S202/T205 (**Figure 7B**). These results demonstrate that the modification pattern and profile of nano-aggregates evolve with the progression of AD stages. This trend was similarly observed for intermediate-aggregates, where there was a greater heterogeneity in the distribution of enrichment scores in AD Braak stage IV compared to AD Braak stage VI, and the percentage of dually modified intermediate-aggregates significantly increased in advanced AD (**Figure 7A, B**). Micro-aggregates, on the other hand, were predominantly dually modified with both combinations of hyperphosphorylated residues visualized in both AD Braak stages IV and VI. These findings suggest that as tau aggregation advances, the aggregates accumulate more tau that is modified at different residues, either through continued phosphorylation or via co-aggregation of differentially modified tau seeds.

**Figure 7:**
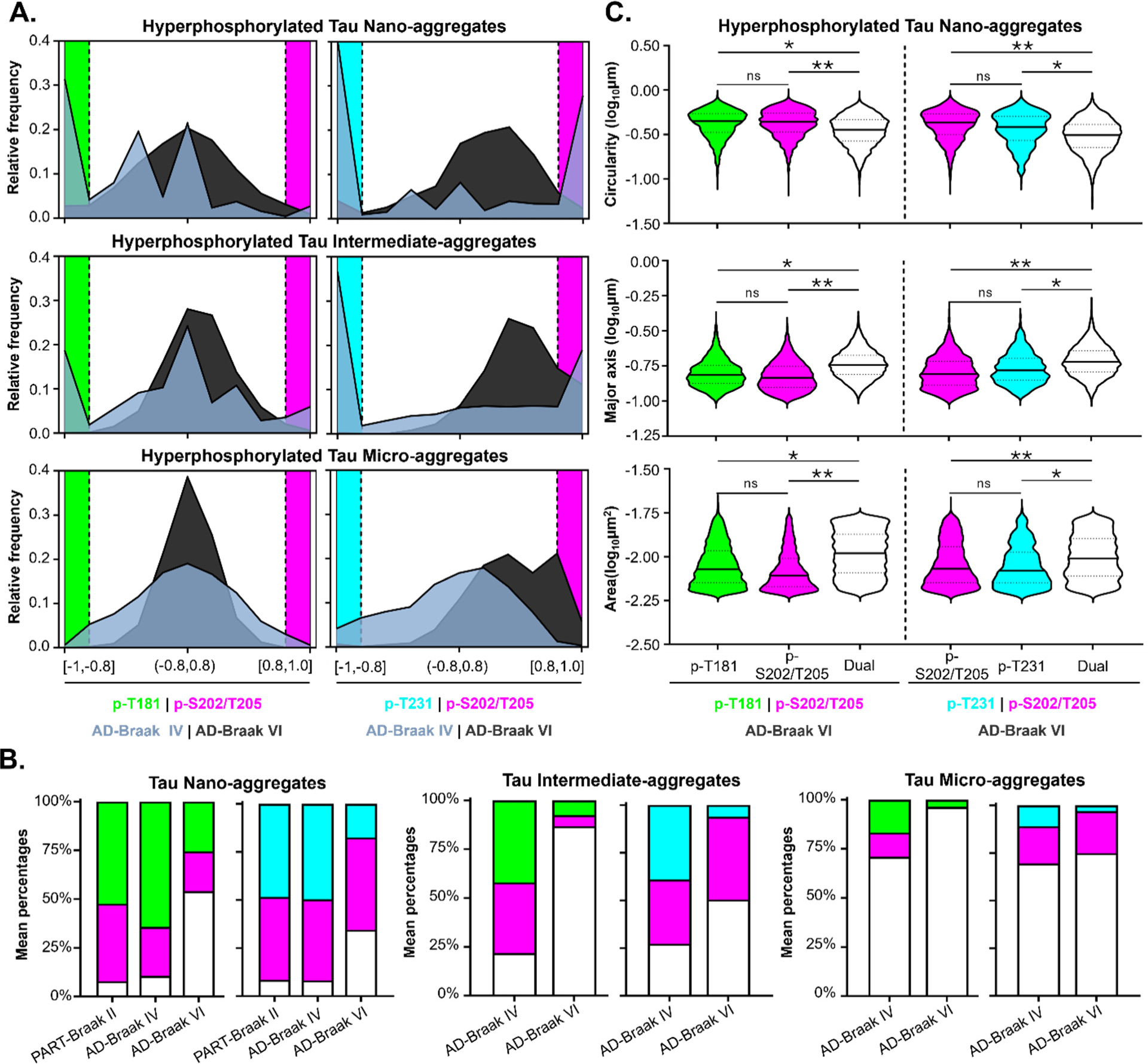
Nano-aggregates are differentially hyperphosphorylated at advanced AD stages. **(A)** Histograms showing the enrichment score of classified tau aggregates from AD-Braak IV (blue grey) and VI (dark grey) immunolabeled with phospho-specific primary antibody combinations: AT8 (magenta) + AT270 (green); AT8 (magenta) + AT180 (cyan), to identify p-S202/T205, p-T181, and p-T231, respectively. Enrichments score of −0.8 to −1 (shaded green or shaded cyan) and 0.8 to 1 (shaded magenta) were taken to correspond to singly hyperphosphorylated tau aggregates. Enrichment scores between −0.8 to 0.8 were taken to correspond to dually hyperphosphorylated tau aggregates. AT8 + AT270 immunolabeled samples were imaged with combination of imager-probes 1 and 2 (ATTO655,Cy3b); AT8 + AT180 immunolabeled samples were imaged with combination of imager-probes 1 and 2 (ATTO655,Cy3b). **(B)** Percentage of dually hyperphosphorylated (white) and singly hyperphosphorylated (magenta for p-S202/T205, cyan for p-T231 and green for p-T181) tau nano- (left), intermediate- (middle) and micro-aggregates (right). Stacked bar plots show the mean percentage. Magenta = p-S202/T205, Green = p-T181, Cyan = p-T231, White= dual-modified. PART-Braak II: N = 4 tissue sections, n = 16 fields of view; AD-Braak IV: N = 4 tissue sections, n = 16 fields of view; AD-Braak VI: N = 4 tissue sections, n = 16 fields of view. **(C)** Violin plots showing the circularity, major axis, and area for p-T181 only (green), p-S202/T205 only (magenta), p-T231 only (cyan), or p-T181/p-S202/T205 and p-T231/p-S202/T205 dually modified (white) hyperphosphorylated tau nano-aggregates in AD-Braak VI. Solid line indicates median, and the dotted lines indicate the 25th and 75th percentile. AD-Braak VI: N = 4 tissue sections, n = 16 fields of view. A Wilcoxon rank sum test was performed between enrichment score specific populations of tau aggregates in AD-stage VI. A p value of < 0.05 was taken as statically significant. P values = (ns) >0.05, (*) 0.05 – 0.01, (**) < 0.01.

Given the changes in the phosphorylation patterns of tau nano-aggregates across different AD-Braak stages, we sought to determine whether dually modified nano-aggregates differed morphologically from their singly modified counterparts. ECLiPSE analysis revealed that dually modified nano-aggregates exhibited reduced circularity, increased major axis length, and larger area compared to the singly modified nano-aggregates (**Figure 7C, Supplementary Figure 6A,B**), resembling the morphological characteristics that distinguish nano-aggregates in AD from those observed in PART.

Overall, these results reveal that the phosphorylation patterns of tau aggregates evolve as the aggregation progresses and that dually modified nano-aggregates display distinct morphological features compared to their singly modified counterparts.

## Discussion

Utilizing DNA-PAINT super-resolution microscopy, we demonstrate the feasibility of detecting hyperphosphorylated nano-sized tau aggregates (nano-aggregates) within human postmortem brain tissues corresponding to PART and various AD stages. While the specific tau protein copy number and biochemical attributes (e.g., soluble tau oligomers versus insoluble tau aggregates) of these tau nano-aggregates remain elusive, they likely correspond to tau seeds such as tau oligomers and small fibrils, collectively representing the early phases of tau aggregation. Previously, in engineered cell models, we employed calibration approaches to quantify the protein copy number of nanoscale tau aggregates and found that they correspond to tau oligomers containing more than 4 tau proteins (*63*). The punctate tau nano-aggregates identified in PART and AD are presumed to be tau oligomers, given their morphological similarity to those observed in engineered cell models. Future work, using oligomer- and conformation-specific antibodies alongside calibration and quantification approaches, will further elucidate the properties of tau nano-aggregates.

Importantly, our results underscore the prevalence of nano-aggregates as the predominant species of tau aggregates in PART and AD, highlighting the sensitivity of DNA-PAINT in detecting and quantifying their presence. In addition, leveraging the high resolution and quantitative nature of DNA-PAINT, we demonstrated that nano-aggregates exhibit distinct morphological characteristics between PART and AD. This suggests that the nature of these aggregates may be specific to the disease and its Braak stage, and those observed in PART may even represent more physiological tau nano-aggregates. In the future, it would be interesting to compare the morphology of tau nano-aggregates between young and aged brains or across different tauopathies to determine whether each tauopathy is marked by morphologically unique tau nano-aggregates. Our results also show that nano-aggregates contain tau hyperphosphorylated at many disease relevant residues (p-T231, p-T181 and pS202/T205) found within tau’s PRR region. We focused on these particular phospho-tau residues due to their significance in disease progression, their link to specific stages of tau aggregation, and the availability of highly specific and thoroughly validated antibodies (*68*). Looking ahead, it would be worthwhile to explore other post-translational modifications of tau, including other phospho-tau residues such as Threonine 217 (another tau biomarker), and ubiquitination marks, the latter of which has been shown biochemically to be enriched in tau oligomers (*71*).

Multi-color DNA-PAINT analysis revealed that tau nano-aggregates in advanced AD possess a combination of phospho-tau residues, such as both p-T181 and p-S202/T205 or p-T231 and p-S202/T205. This finding aligns with a recent study that employed single-molecule pull-down techniques to investigate the phospho-tau composition of small tau aggregates extracted from human brain tissues and biofluids (*72*). Given the species overlap between the antibodies, we could not simultaneously examine all three phospho-tau modifications in a multiplexed approach. Nonetheless, it is probable that a subset of the nano-aggregates carries all three phospho-tau modifications. In the future, directly conjugating primary antibodies with DNA-PAINT compatible docking oligos will facilitate the simultaneous examination of a broader range of tau’s post-translational modifications to determine the combinatorial modification signature unique to different tau aggregates.

In contrast to advanced AD, nano-aggregates from cases of intermediate AD and PART mainly contained only one of the phospho-tau modifications visualized. This result is surprising especially considering that p-T181— one of the modifications we visualized —has previously been described as a “master” residue that, once phosphorylated, can lead to further phosphorylation at other residues (*36*). Our results suggest that initial formation of nano-aggregates likely does not require a combination of tau modifications, although we cannot exclude that other phospho-tau modifications or PTMs may be enriched in these nano-aggregates. However, as the disease progresses and aggregation continues, tau aggregates may undergo further hyperphosphorylation. It is also possible that nano-aggregates with different modifications could coalesce, forming aggregates with combined modifications. Supporting this possibility, our analysis showed that nano-aggregates with dual modifications were morphologically different from those with a single modification, often being larger and less circular. These combinatorically modified and morphologically distinct nano-aggregates might exhibit greater toxicity and a higher propensity to seed further aggregation, particularly given their prevalence in advanced AD and the observation that micro-aggregates are often dually modified. Future work comparing different tauopathies at various disease stages and utilizing markers for several disease-relevant tau PTMs has the potential to reveal the specific modifications and morphological traits of tau seeds. This information may provide crucial insights into the critical modifications to target for inhibiting tau aggregation.

Overall, the improved capability to detect and quantitatively analyze tau nano-aggregates in postmortem human brain tissue paves the way for new research into the molecular mechanisms of Alzheimer’s Disease (AD) and related tauopathies.

## Materials and Methods

### Tissue Samples

Formalin-fixed and paraffin embedded temporal cortex tissue blocks from three cases neuropathologically diagnosed with PART (Primary Age-Related Tauopathy), Intermediate Alzheimer’s disease, and Advanced Alzheimer’s disease were obtained from the University of Pennsylvania (Penn) Center for Neurodegenerative Disease Research (CNDR) Center as outlined in Table 1. Each tissue block was cut into 6 μm thick sections using a microtome and mounted over Poly-L-Lysine coated coverslips. Mounted sections were incubated at 37°C overnight and stored at room temperature.

### Tissue preparation

Mounted tissue sections were incubated with Xylene (99% vol/vol; Sigma Aldrich) two times (5 minutes per incubation). Samples were used right away after deparaffinization and not stored. Deparaffinized samples were incubated with Ethanol two times (1 minute per incubation). Then samples were rehydrated by incubating them in a series of Ethanol and deionized water solutions of 90%, 80%, and 70% (vol/vol; Thermo Fisher) (1 minute per incubation). Lastly, samples were quickly rinsed with deionized water three times.

### Immunolabeling

The immunolabeling consisted of six main steps: antigen retrieval, reduction of autofluorescence, permeabilization, blocking, post-fixation, and clearing (*73,74*). For antigen retrieval, freshly deparaffinized and rehydrated samples were incubated with Tris-EDTA buffer solution (10mM Tris Base, 1mM EDTA, 0.05% Tween-20, PBS, pH 9.0) for 15 minutes at 94°C and washed three times with PBS (5 min per wash). For reduction of autofluorescence, samples were incubated with freshly made 0.1% (wt/vol) sodium borohydride in PBS solution on ice for 15 minutes and washed three times with PBS (5 min per wash). Then, samples were permeabilized with 0.2% Triton x-100 (vol/vol; Sigma-Aldrich) in PBS for 30 minutes and washed three times with PBS (5 min per wash). Next, samples were blocked for 1 hour with blocking solution (3% (wt/vol) BSA, 0.2% Triton, PBS) at room temperature and washed three times with PBS (5 min per wash). They were then incubated overnight at 4°C in a humidified chamber with the appropriate dilution of primary antibody in blocking solution and washed three times with PBS (5 min per wash). Samples were then incubated for 2 hours at room temperature with a dilution of the appropriate secondary antibody and washed with PBS (5 min per wash). Then, samples were post-fixed with 4% PFA in PBS for 10 minutes at room temperature and washed three times with PBS (5 min per wash). Lastly, samples were treated with 60% 2,2-thiodiethanol (vol/vol; Sigma Aldrich) in PBS for 30 minutes at room temperature to reduce background and optical aberrations.

For negative control samples, all the above steps were followed except for the addition of primary antibody.

Labeling of the samples for dual-color imaging was carried out sequentially. Samples were first incubated with one of the primary antibodies using the protocol just described. After the incubation step with secondary antibody, samples were post-fixed with 4% PFA in PBS for 10 minutes at room temperature and washed three times with PBS (5 min per wash). Samples were then blocked for 30 minutes with blocking solution (3% (wt/vol) BSA, 0.2% Triton, PBS) at room temperature and washed three times with PBS (5 min per wash). The samples were then incubated overnight at 4°C in a humidified chamber with the appropriate dilution of the second primary antibody in blocking solution and washed three times with PBS (5 min per wash). Afterwards, samples were incubated for 2 hours at room temperature with a dilution of the appropriate secondary antibody and washed three times with PBS (5 min per wash). Then, samples were post-fixed once again with 4% PFA in PBS for 10 minutes at room temperature and washed with PBS (5 min per wash). Lastly, samples were treated with 60% TDE (vol/vol; Sigma Aldrich) in PBS for 30 minutes at room temperature.

A list of primary and secondary antibodies used in these studies is provided in Tables 2 and 3. The primary antibodies we used for single-color acquisitions were: AT8 at a 1:50 dilution, AT180 at a 1:100 dilution, and AT270 at a 1:50 dilution. The secondary antibodies we used in these studies were at a 1:100 dilution.

### Confocal Imaging

For confocal imaging, all of the above steps were followed except with the addition of a DAPI stain, and different dilutions of primary and secondary antibodies. After the clearing step, samples were incubated with a (1:5000) DAPI solution (vol/vol) for 10 minutes then washed three times with PBS (5 min per wash). The primary antibodies used were: AT8 at a 1:200 dilution, AT180 at a 1:400 dilution, and AT270 at a 1:200 dilution. The secondary antibodies we used for these images were at a 1:500 dilution. AT8 labeled samples were imaged with donkey anti-mouse Alexa-647 (secondary antibody), and AT180 and AT270 labeled samples were imaged with Donkey anti-rabbit Alexa-546 (secondary antibody).

All confocal images were acquired using Zeiss LSM 980 with Airyscan 2 inverted confocal microscope, with a Plan-Apochromat 20x/0.8 M27 objective. Z-stacks were acquired in frame mode, at scan speed 9 and using an average of 4. Image size was 1536×1536 pixels by 10-12 Z-stacks, with voxel size 0.28×0.28×1 microns (x-y-z). All fields of view included in this manuscript were randomly selected and imaged. Maximum intensity projections were generated using ImageJ/Fiji (NIH) (*75*).

We acknowledge the Cell & Developmental Biology (CDB) Microscopy Core Facility in the Perelman School of Medicine at the University of Pennsylvania for the Zeiss LSM 980 imaging.

### DNA-PAINT Imaging

All DNA-PAINT images were acquired using the Oxford Nanoimaging System. All fields of view included in this manuscript were randomly selected and imaged. The Nanoimager-S microscope has the following configuration: 405-, 488-, 561-, and 640-nm lasers, 498-551- and 576-620-nm band-pass filters in channel 1, and 666-705-nm band-pass filters in channel 2, 100x 1.45 NA oil immersion objective (Olympus), and Hamamatsu Flash 4 V3 sCMOS camera. All single-and dual-color acquisitions were conducted at 27°C using HiLo illumination.

For single-color DNA-PAINT acquisitions, samples and negative controls were incubated with a solution of appropriate imager-probe in imaging buffer (0.5nM; vol/vol). AT8 labeled samples were imaged with imager-probe 1 (ATTO655), and AT180 and AT270 labeled samples were imaged with imager-probe 2 (Cy3b). Negative control samples labeled with anti-mouse docking site 1 (secondary antibody) were imaged with imager-probe 1 (ATTO655), whereas negative samples labeled with anti-rabbit docking site 2 (secondary antibody) were imaged with imager-probe 2 (Cy3b). All images were acquired at a 100 ms exposure for 34,000 frames. The imager-probe solution was changed every 4 images (∼ every 4 hours). Prior to the addition of a fresh imager-probe solution, samples were thoroughly washed 5 times with PBS (3 min per wash).

For dual-color DNA-PAINT acquisitions, samples were incubated with a solution of appropriate imager-probes (0.5nM; vol/vol) in imaging buffer. AT8, AT180 dually labeled samples were imaged with solution of imager-probes 1 and 2 (ATTO655, Cy3b), and AT8,AT270 dually labeled samples were imaged with solution of imager-probes 1 and 2 (ATTO655, Cy3b). Dual-color images were acquired at a 100 ms exposure using a laser program in which the 561- and 647-lasers were sequentially activated every 200 frames, capturing a total of 68,000 frames, 34,000 frames per target. The solution of imager-probes was changed every 2 images (∼ every 4 hours). Prior to the addition of a fresh imager-probe solution, samples were thoroughly washed five times with PBS (3 min per wash).

### Data Analysis

#### A. Single-Color DNA-PAINT Images

##### Voronoi Tessellation and Segmentation

For single-color acquisitions, localizations were exported in .csv files using the NimOS localization software and rendered using a custom-written data analysis software (https://github.com/melikelakadamyali/StormAnalysisSoftware; MATLAB R2022a). All subsequent data analysis were performed using this custom-written software. For data segmentation, we first performed Voronoi tessellation, in which each localization in space is associated to a Voronoi polygon based on its neighboring localizations (*69*). The Voronoi polygon area is a measure of localization density, with dense localizations having small Voronoi polygons areas and vice versa. The Voronoi polygon area can therefore be used as a threshold to cluster together localizations in dense regions for segmentation (*69*). The area threshold for images from each case were adjusted accordingly to account for differences in tau aggregate burden and localization density: PART-Braak II = (0.00018 – 0.0003 μm^2^); AD-Braak IV = (0.001 – 0.002 μm^2^); AD-Braak VI = (>0.002 μm^2^). Additionally, we employed a threshold of minimum 10 localizations, in which segmented objects having less than 10 localizations were discarded. We visually confirmed that the selected thresholds properly segmented individual tau aggregates in the single-color super-resolution images. These thresholds were applied consistently across all imaged brain tissues.

##### Removal of Imaging Artefacts

Segmented objects have a positive correlation between the area value and number of localizations. Objects that deviate from this trend (i.e. that have a small area value and a large number of localizations, or large area values and a small number of localizations) correspond to imaging artefacts (*63*). These outlier objects were further filtered out from our lists of segmented tau aggregates for all images.

##### Removal of Background and Noise

Images of negative control samples were segmented using the same Voronoi polygon area threshold and minimum number of localizations as described above. The area distribution of the resulting segmented objects was plotted and compared to the area distribution of objects from the corresponding positive control samples. An area threshold was then imposed based on this comparison to remove additional background and noise. For samples imaged with the imager-probe 2 (Cy3b), the area threshold was 0.004 μm^2^, and for those imaged with imager-probe 1 (ATTO655), the area threshold was 0.006 μm^2^. The area thresholds were slightly different for the two imager-probes as they gave rise to slightly different levels of background/noise. Applying these thresholds to the negative control samples removed >97% of segmented objects corresponding to background and noise. Hence, these thresholds were used for removing background and noise present in the positive control samples imaged using the corresponding imager-probes.

##### Area-based Classification of Segmented Tau Aggregates

To separate segmented tau aggregates into distinct classes, we evaluated the area distribution of tau aggregates across cases and established specific area thresholds for separating segmented tau aggregates into three distinct area-based classes (Class I: nano-aggregates, Class II: intermediate-sized aggregates, Class III: micro-sized aggregates). The area thresholds used were 0.004 – 0.017 μm^2^ (for aggregates visualized using imager-probe 2 (Cy3b)) or 0.006-0.017 μm^2^ (for aggregates visualized using imager-probe 1 (ATTO655)) for Class I, 0.017 – 0.15 μm^2^ for Class II, and > 0.15 μm^2^ for Class III. Note that the minimum area threshold for Class I was slightly different for the nano-aggregates imaged using the two different imager-probes as described above. These thresholds were applied consistently across all images. We visually confirmed that tau aggregates from single-color acquisitions from all cases were properly classified.

##### ECLiPSE Analysis

ECLiPSE is an automated shape classification method that is capable of determining morphological properties of structures imaged using super-resolution microscopy (*67*). For detailed information on the ECLiPSE methodology, we refer the reader to (*67*). Briefly, the localized point clouds were used, without any additional rendering, to calculate 67 different morphological and structural descriptors (i.e., geometric, boundary, skeleton, fractal, etc. descriptors). The use of the unrendered point cloud data provides the most unbiased quantification of the morphological properties.

Once all properties were calculated with ECLiPSE, Principal Component Analysis (PCA) was used to capture the essential information in the data (after unit variance scaling to remove scaling artifacts). PCA is a compression technique that transforms data into latent variables (called principal components, PC) by using a linear combination of the original variables with the goal of maximizing data variability. These latent variables are uncorrelated with respect to each other and will create a new coordinate system onto which the original data can be projected. As each latent variable explains more data variability than the subsequent one, the first few PCs retain most of the variability of the original data set. This allows us to simplify complex data sets (i.e. dimensionality reduction) while retaining the essential characteristics of the data. This property of PCA also explains that data points that are spatially close in the PCA space will have similar features (i.e., biological properties using ECLiPSE), and therefore, two sets of data points that overlap will have similar biological properties, whereas two sets of data points that do not overlap will have different biological properties. After constructing the PCA space and projecting the data points into this new coordinate system, each data point was color coded according to which case it belongs (PART-Braak II, AD-Braak IV, or AD-Braak VI).

##### Statistical Analysis

All unpaired, non-parametric Mann-Whitney tests performed in these studies were calculated using GraphPad Prism, version 9.4.1. A p value of < 0.05 was taken as statically significant. P values = (ns) >0.05, (*) 0.05 – 0.03, (**) 0.002 – 0.03, (***) 0.0002 – 0.002, (****) 0.0001 – 0.0002. All Wilcoxon rank sum tests were performed using MATLAB 2022b. A p value of < 0.05 was taken as statically significant. P values = (ns) >0.05, (*) 0.05 – 0.01, (**) < 0.01.

#### B. Dual-color DNA-PAINT Images

##### Voronoi Tessellation and Segmentation

For dual-color acquisitions, a strategy similar to that of single-color acquisitions was applied, with the exception that prior to Voronoi tessellation, channels pertaining to the localizations from each target were combined to generate a merged reference image. Segmentation was then performed as described above followed by the removal of imaging artifacts and classification of segmented tau aggregates into area-based classes.

To correct for segmentation errors originating from the merging of channels, the segmentation quality was evaluated by calculating the overlap score for the localizations belonging to the two distinct channels within an individual aggregate. The overlap score was computed by first transforming the localizations of each channel into an alpha-shape (i.e., a special case of Voronoi polygons that is a precise description of the aggregates) and then determining the intersection between these two alpha-shape objects. The overlap score is then calculated as the percentage of area that overlaps (i.e., the area of this intersection) with the alpha-shape object of the localizations in the other channel. This was evaluated with respect to both channels and the overlap scores were consistent, regardless of which channel was used as a reference. We considered segmented aggregates with an overlap score <30% to be two distinct aggregates that were spatially-proximate and became merged in the reference image in error. To rectify such segmentation errors, we first applied an area filter (above 0.15 μm^2^) to remove all micron-sized aggregates as these had an overlap score greater than 30% and did not need to be corrected. Aggregates below this area threshold with an overlap score of <30% were re-segmented. These now distinct aggregates were also reassigned to the correct area-based class (i.e., nano-, intermediate-, or micro-aggregate) if needed. Resulting lists of segmented aggregates were combined and visually inspected for proper segmentation once again. These thresholds and re-segmentation parameters were applied consistently across different conditions.

##### Colocalization Analysis

Additionally, to evaluate the composition of the different aggregates, an enrichment score was calculated. This enrichment score is obtained by calculating the ratio between the difference in number of localizations in each channel of the aggregate and the total number of localizations for the aggregate. An enrichment score of −1 or 1 means that the aggregate completely consists of a single type of phospho-tau marker, whereas an enrichment score of 0 represents an equal distribution of phospho-tau markers for that aggregate. Anything in between −1 and 0, or 0 and 1 will contain more of one phospho-tau marker than the other.

## Acknowledgments

We acknowledge the Cell & Developmental Biology (CDB) Microscopy Core Facility in the Perelman School of Medicine at the University of Pennsylvania and Dr. Natali Chanaday for help with confocal imaging. We acknowledge Bilan Yakoub for help with preliminary sample preparation and imaging at the early stages of this project.

## Author Contributions

Conceptualization: ML and ANSR

Methodology: ANSR, SH, CB

Investigation: ANSR, SH

Analysis: ANSR, SH

Visualization: ANSR

Materials: EBL

Supervision: ML, EBL

Writing – original draft: ML, ANSR

Writing – review & editing: ML, ANSR, SH, CB, EBL

## Data and materials availability

The code used in this manuscript has been deposited to Figshare (Doi: 10.6084/m9.figshare.25673916). The processed and analyzed data has been deposited to https://github.com/melikelakadamyali/StormAnalysisSoftware. ECLIPSE software has been deposited to https://github.com/LakGroup/ECLiPSE. The raw imaging data is too large to deposit to a public repository and is available from authors upon request. All other data are available in the main text and supplementary materials.

## References

1. T. Guo, W. Noble, D. P. Hanger, Roles of tau protein in health and disease. Acta Neuropathol 133, 665–704 (2017).

2. V. M. Lee, M. Goedert, J. Q. Trojanowski, Neurodegenerative tauopathies. Annu Rev Neurosci 24, 1121–1159 (2001).

3. Y. Wang, E. Mandelkow, Tau in physiology and pathology. Nature Reviews Neuroscience 17, 22–35 (2016).

4. A. Cario, C. L. Berger, Tau, microtubule dynamics, and axonal transport: New paradigms for neurodegenerative disease. Bioessays 45, e2200138 (2023).

5. L. Balabanian et al., Tau differentially regulates the transport of early endosomes and lysosomes. Mol Biol Cell 33, ar128 (2022).

6. J. L. Stern, D. V. Lessard, G. J. Hoeprich, G. A. Morfini, C. L. Berger, Phosphoregulation of Tau modulates inhibition of kinesin-1 motility. Mol Biol Cell 28, 1079–1087 (2017).

7. R. Dixit, J. L. Ross, Y. E. Goldman, E. L. Holzbaur, Differential regulation of dynein and kinesin motor proteins by tau. Science 319, 1086–1089 (2008).

8. N. Samudra, C. Lane-Donovan, L. VandeVrede, A. L. Boxer, Tau pathology in neurodegenerative disease: disease mechanisms and therapeutic avenues. J Clin Invest 133, (2023).

9. Y. Zhang, K.-M. Wu, L. Yang, Q. Dong, J.-T. Yu, Tauopathies: new perspectives and challenges. Molecular Neurodegeneration 17, 28 (2022).

10. J. Gotz, G. Halliday, R. M. Nisbet, Molecular Pathogenesis of the Tauopathies. Annu Rev Pathol 14, 239–261 (2019).

11. M. G. Spillantini, M. Goedert, Tau pathology and neurodegeneration. Lancet Neurol 12, 609–622 (2013).

12. L. I. Binder, A. L. Guillozet-Bongaarts, F. Garcia-Sierra, R. W. Berry, Tau, tangles, and Alzheimer’s disease. Biochim Biophys Acta 1739, 216–223 (2005).

13. H. Braak, E. Braak, Neuropathological stageing of Alzheimer-related changes. Acta Neuropathol 82, 239–259 (1991).

14. T. J. Montine et al., National Institute on Aging–Alzheimer’s Association guidelines for the neuropathologic assessment of Alzheimer’s disease: a practical approach. Acta Neuropathologica 123, 1–11 (2012).

15. J. A. Trejo-Lopez, A. T. Yachnis, S. Prokop, Neuropathology of Alzheimer’s Disease. Neurotherapeutics 19, 173–185 (2022).

16. T. Vogels et al., Propagation of Tau Pathology: Integrating Insights From Postmortem and In Vivo Studies. Biol Psychiatry 87, 808–818 (2020).

17. N. Franzmeier et al., Functional brain architecture is associated with the rate of tau accumulation in Alzheimer’s disease. Nature Communications 11, 347 (2020).

18. L. Martin, X. Latypova, F. Terro, Post-translational modifications of tau protein: implications for Alzheimer’s disease. Neurochem Int 58, 458–471 (2011).

19. C. Alquezar, S. Arya, A. W. Kao, Tau Post-translational Modifications: Dynamic Transformers of Tau Function, Degradation, and Aggregation. Front Neurol 11, 595532 (2020).

20. G. Simic et al., Tau Protein Hyperphosphorylation and Aggregation in Alzheimer’s Disease and Other Tauopathies, and Possible Neuroprotective Strategies. Biomolecules 6, 6 (2016).

21. C. X. Gong, K. Iqbal, Hyperphosphorylation of microtubule-associated protein tau: a promising therapeutic target for Alzheimer disease. Curr Med Chem 15, 2321–2328 (2008).

22. E. Kopke et al., Microtubule-associated protein tau. Abnormal phosphorylation of a non-paired helical filament pool in Alzheimer disease. J Biol Chem 268, 24374–24384 (1993).

23. E. S. Matsuo et al., Biopsy-derived adult human brain tau is phosphorylated at many of the same sites as Alzheimer’s disease paired helical filament tau. Neuron 13, 989–1002 (1994).

24. F. Liu et al., Site-specific effects of tau phosphorylation on its microtubule assembly activity and self-aggregation. Eur J Neurosci 26, 3429–3436 (2007).

25. E. Ercan et al., A validated antibody panel for the characterization of tau post-translational modifications. Molecular Neurodegeneration 12, 87 (2017).

26. M. Goedert, R. Jakes, E. Vanmechelen, Monoclonal antibody AT8 recognises tau protein phosphorylated at both serine 202 and threonine 205. Neurosci Lett 189, 167–169 (1995).

27. J. C. Augustinack, A. Schneider, E. M. Mandelkow, B. T. Hyman, Specific tau phosphorylation sites correlate with severity of neuronal cytopathology in Alzheimer’s disease. Acta Neuropathol 103, 26–35 (2002).

28. J. Neddens et al., Phosphorylation of different tau sites during progression of Alzheimer’s disease. Acta Neuropathologica Communications 6, 52 (2018).

29. I. Alafuzoff et al., Staging of neurofibrillary pathology in Alzheimer’s disease: a study of the BrainNet Europe Consortium. Brain Pathol 18, 484–496 (2008).

30. L. Amniai, G. Lippens, I. Landrieu, Characterization of the AT180 epitope of phosphorylated Tau protein by a combined nuclear magnetic resonance and fluorescence spectroscopy approach. Biochem Biophys Res Commun 412, 743–746 (2011).

31. G. A. Jicha et al., A conformation- and phosphorylation-dependent antibody recognizing the paired helical filaments of Alzheimer’s disease. J Neurochem 69, 2087–2095 (1997).

32. C. Schaffer et al., Biomarkers in the Diagnosis and Prognosis of Alzheimer’s Disease. J Lab Autom 20, 589–600 (2015).

33. M. Suarez-Calvet et al., Novel tau biomarkers phosphorylated at T181, T217 or T231 rise in the initial stages of the preclinical Alzheimer’s continuum when only subtle changes in Abeta pathology are detected. EMBO Mol Med 12, e12921 (2020).

34. K. L. Meeker et al., Comparison of cerebrospinal fluid, plasma and neuroimaging biomarker utility in Alzheimer’s disease. Brain Commun 6, fcae081 (2024).

35. V. Papaliagkas et al., CSF Biomarkers in the Early Diagnosis of Mild Cognitive Impairment and Alzheimer’s Disease. Int J Mol Sci 24, (2023).

36. K. Stefanoska et al., Alzheimer’s disease: Ablating single master site abolishes tau hyperphosphorylation. Sci Adv 8, eabl8809 (2022).

37. Y. Carlomagno et al., The AD tau core spontaneously self-assembles and recruits full-length tau to filaments. Cell Rep 34, 108843 (2021).

38. M. Usenovic et al., Internalized Tau Oligomers Cause Neurodegeneration by Inducing Accumulation of Pathogenic Tau in Human Neurons Derived from Induced Pluripotent Stem Cells. J Neurosci 35, 14234–14250 (2015).

39. C. Kim et al., Distinct populations of highly potent TAU seed conformers in rapidly progressing Alzheimer’s disease. Sci Transl Med 14, eabg0253 (2022).

40. C. A. Lasagna-Reeves et al., Identification of oligomers at early stages of tau aggregation in Alzheimer’s disease. FASEB J 26, 1946–1959 (2012).

41. S. Boluda et al., Differential induction and spread of tau pathology in young PS19 tau transgenic mice following intracerebral injections of pathological tau from Alzheimer’s disease or corticobasal degeneration brains. Acta Neuropathol 129, 221–237 (2015).

42. S. Narasimhan et al., Pathological Tau Strains from Human Brains Recapitulate the Diversity of Tauopathies in Nontransgenic Mouse Brain. J Neurosci 37, 11406–11423 (2017).

43. S. K. Kaufman et al., Tau Prion Strains Dictate Patterns of Cell Pathology, Progression Rate, and Regional Vulnerability In Vivo. Neuron 92, 796–812 (2016).

44. J. L. Guo et al., Unique pathological tau conformers from Alzheimer’s brains transmit tau pathology in nontransgenic mice. J Exp Med 213, 2635–2654 (2016).

45. F. Clavaguera et al., Brain homogenates from human tauopathies induce tau inclusions in mouse brain. Proc Natl Acad Sci U S A 110, 9535–9540 (2013).

46. A. Van der Jeugd et al., Cognitive defects are reversible in inducible mice expressing pro-aggregant full-length human Tau. Acta Neuropathol 123, 787–805 (2012).

47. C. A. Lasagna-Reeves et al., Tau oligomers impair memory and induce synaptic and mitochondrial dysfunction in wild-type mice. Molecular Neurodegeneration 6, 39 (2011).

48. D. L. Castillo-Carranza et al., Specific targeting of tau oligomers in Htau mice prevents cognitive impairment and tau toxicity following injection with brain-derived tau oligomeric seeds. J Alzheimers Dis 40 Suppl 1, S97–S111 (2014).

49. S. L. DeVos et al., Synaptic Tau Seeding Precedes Tau Pathology in Human Alzheimer’s Disease Brain. Front Neurosci 12, 267 (2018).

50. D. W. Sanders et al., Distinct tau prion strains propagate in cells and mice and define different tauopathies. Neuron 82, 1271–1288 (2014).

51. F. Clavaguera et al., “Prion-like” templated misfolding in tauopathies. Brain Pathol 23, 342–349 (2013).

52. J. L. Guo, V. M. Lee, Seeding of normal Tau by pathological Tau conformers drives pathogenesis of Alzheimer-like tangles. J Biol Chem 286, 15317–15331 (2011).

53. S. S. Shafiei, M. J. Guerrero-Munoz, D. L. Castillo-Carranza, Tau Oligomers: Cytotoxicity, Propagation, and Mitochondrial Damage. Front Aging Neurosci 9, 83 (2017).

54. X. Q. Chen, W. C. Mobley, Alzheimer Disease Pathogenesis: Insights From Molecular and Cellular Biology Studies of Oligomeric Abeta and Tau Species. Front Neurosci 13, 659 (2019).

55. J. E. Gerson, D. L. Castillo-Carranza, R. Kayed, Advances in therapeutics for neurodegenerative tauopathies: moving toward the specific targeting of the most toxic tau species. ACS Chem Neurosci 5, 752–769 (2014).

56. E. Ercan-Herbst et al., A post-translational modification signature defines changes in soluble tau correlating with oligomerization in early stage Alzheimer’s disease brain. Acta Neuropathol Commun 7, 192 (2019).

57. S. Dujardin et al., Tau molecular diversity contributes to clinical heterogeneity in Alzheimer’s disease. Nature Medicine 26, 1256–1263 (2020).

58. H. Wesseling et al., Tau PTM Profiles Identify Patient Heterogeneity and Stages of Alzheimer’s Disease. Cell 183, 1699–1713 e1613 (2020).

59. B. Falcon et al., Structures of filaments from Pick’s disease reveal a novel tau protein fold. Nature 561, 137–140 (2018).

60. A. W. P. Fitzpatrick et al., Cryo-EM structures of tau filaments from Alzheimer’s disease. Nature 547, 185–190 (2017).

61. T. Arakhamia et al., Posttranslational Modifications Mediate the Structural Diversity of Tauopathy Strains. Cell 180, 633–644 e612 (2020).

62. C. Bond, A. N. Santiago-Ruiz, Q. Tang, M. Lakadamyali, Technological advances in super-resolution microscopy to study cellular processes. Mol Cell 82, 315–332 (2022).

63. M. T. Gyparaki et al., Tau forms oligomeric complexes on microtubules that are distinct from tau aggregates. Proc Natl Acad Sci U S A 118, (2021).

64. J. L. Guo et al., The Dynamics and Turnover of Tau Aggregates in Cultured Cells: INSIGHTS INTO THERAPIES FOR TAUOPATHIES. J Biol Chem 291, 13175–13193 (2016).

65. R. Jungmann et al., Multiplexed 3D cellular super-resolution imaging with DNA-PAINT and Exchange-PAINT. Nature Methods 11, 313–318 (2014).

66. J. F. Crary et al., Primary age-related tauopathy (PART): a common pathology associated with human aging. Acta Neuropathol 128, 755–766 (2014).

67. S. Hugelier et al., ECLiPSE: A Versatile Classification Technique for Structural and Morphological Analysis of Super-Resolution Microscopy Data. BioRxiv, (2023).

68. M. J. Ellis et al., Validation of Tau Antibodies for Use in Western Blotting and Immunohistochemistry. BioRxiv, (2023).

69. F. Levet et al., SR-Tesseler: a method to segment and quantify localization-based super-resolution microscopy data. Nature Methods 12, 1065–1071 (2015).

70. R. Jungmann et al., Quantitative super-resolution imaging with qPAINT. Nature Methods 13, 439–442 (2016).

71. N. Puangmalai et al., Lysine 63-linked ubiquitination of tau oligomers contributes to the pathogenesis of Alzheimer’s disease. J Biol Chem 298, 101766 (2022).

72. D. Boken et al., Single-Molecule Characterization and Super-Resolution Imaging of Alzheimer’s Disease-Relevant Tau Aggregates in Human Samples. Angew Chem Int Ed Engl, e202317756 (2024).

73. J. Xu et al., Super-resolution imaging reveals the evolution of higher-order chromatin folding in early carcinogenesis. Nature Communications 11, 1899 (2020).

74. C. L. German, M. V. Gudheti, A. E. Fleckenstein, E. M. Jorgensen, in Super-Resolution Microscopy: Methods and Protocols, H. Erfle, Ed. (Springer New York, New York, NY, 2017), pp. 153–162.

75. J. Schindelin et al., Fiji: an open-source platform for biological-image analysis. Nat Methods 9, 676–682 (2012).

