## Supplementary Information for "Super-Resolution Imaging Uncovers Nanoscale Tau Aggregate Hyperphosphorylation Patterns in Human Alzheimer’s Disease Brain Tissue"

Fig. S1.

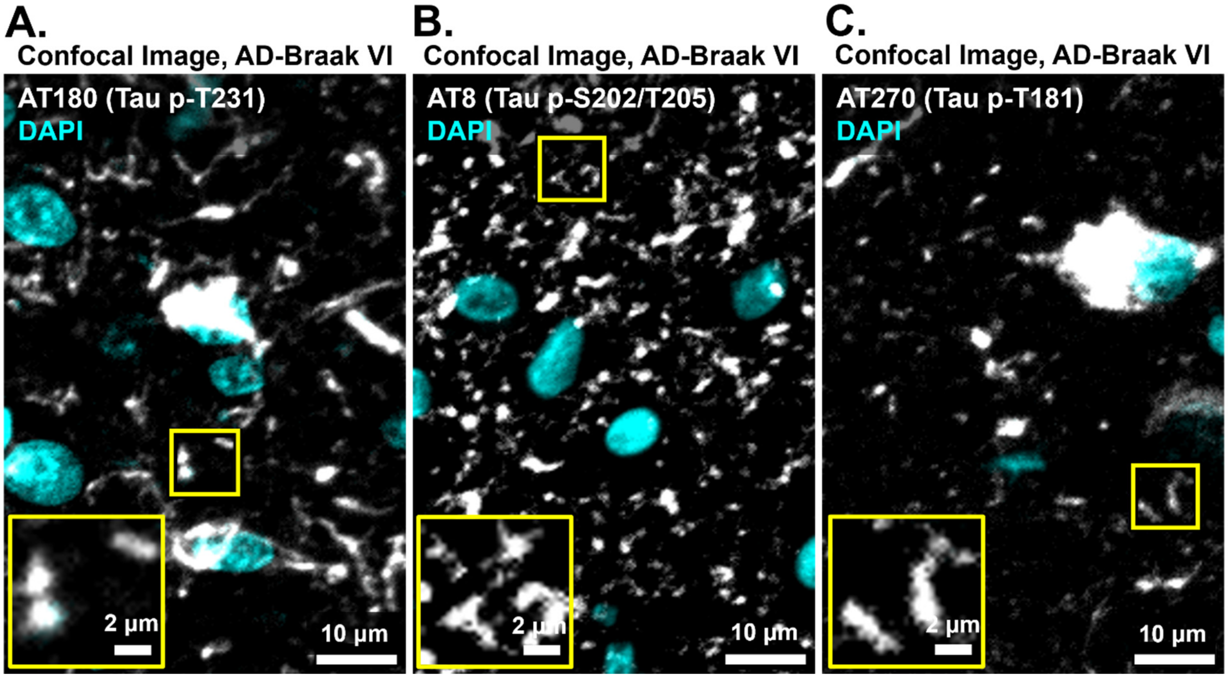


**Single-color confocal images of tau aggregates in AD-Braak VI tissue sections. (A-C)** Representative single-color z-projection confocal image of p-T231 in AD-Braak VI tissue sections immunolabeled with AT180 (white) **(A),** p-S202/T205 immunolabeled with AT8 (white) **(B)**, and p-T181 immunolabeled with AT270 (white) **(C)**. Cyan images show nuclei labeled with DAPI. **(A-C)** AT180 and AT270 labeled samples were imaged with Alexa546 and AT8 labeled samples were imaged with Alexa647.

**Fig. S2.**

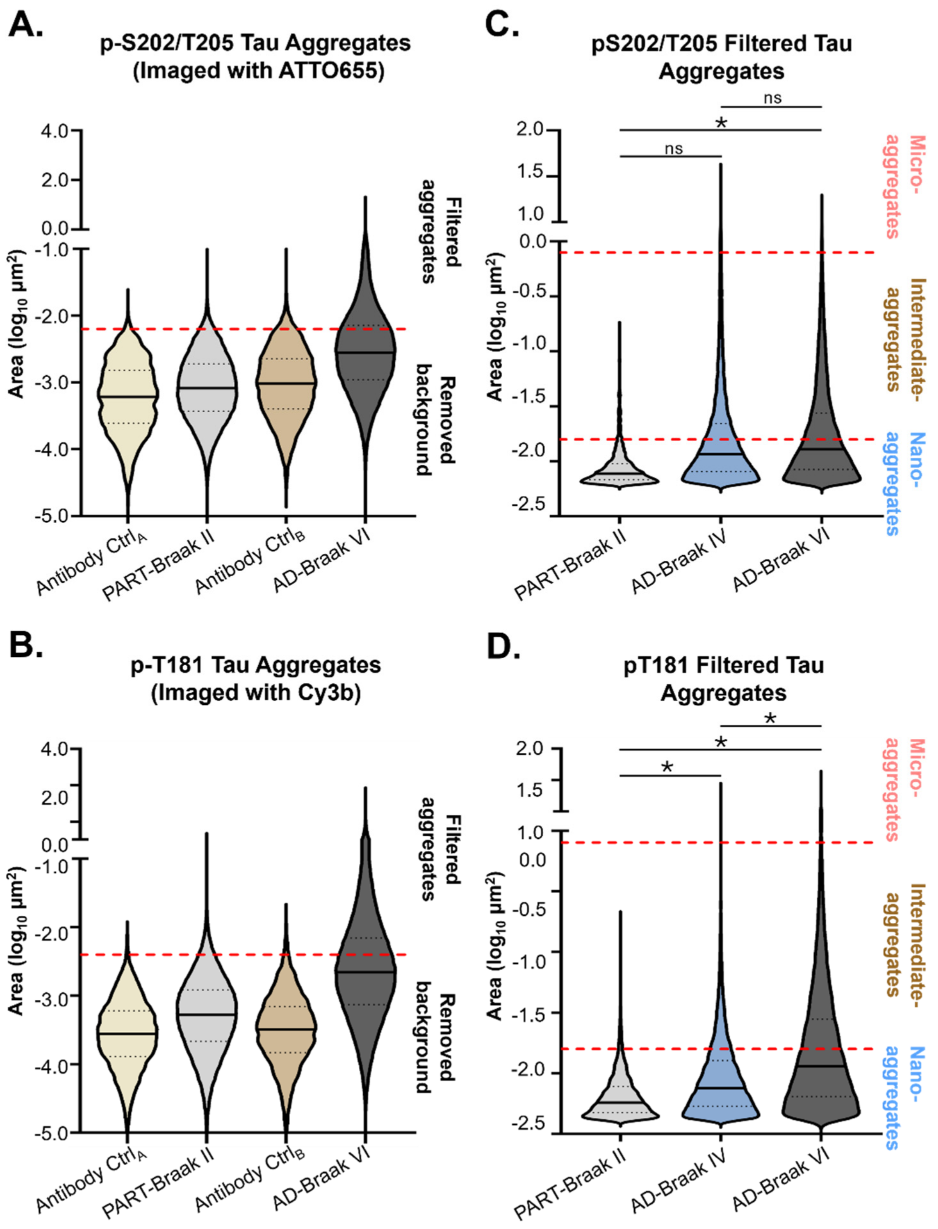


**Negative control experiments for p-S202/T205 imaged with imager-probe 1 (ATTO655) and p-T181 imaged with imager-probe 2 (Cy3b). (A,B)** Violin plots showing the area distribution of segmented objects from the antibody negative control images (Antibody Ctrl$_A$ = PART-Braak II, Antibody Ctrl$_B$ = AD-Braak VI) and positive control samples (PART-Braak II and AD-Braak VI) immunolabeled with AT8 antibody **(A)** or AT270 antibody **(B)**. Red dashed line indicates the area cut-off used for the area-based filtering of background signal. Segmented aggregates below this value are removed from the list of segmented tau aggregates. Solid line indicates median, and the dotted lines indicate the 25th and 75th percentile **(A)** Negative control samples N = 1 tissue section, n = 4 fields of view; positive samples: PART-Braak II: N = 4 tissue sections, n = 17 fields of view; AD-Braak VI: N = 4 tissue sections, n = 18 fields of view. **(B)** Negative control samples N = 1 tissue section, n = 4 fields of view; positive samples: PART-Braak II: N = 4 tissue sections, n = 17 fields of view; AD-Braak VI: N = 4 tissue sections, n = 18 fields of view. **(C, D)** Violin plots showing the area (log10 $\mu m^2$) per tau aggregates after background filtering for p-S202/T205 **(C)** and p-T181 **(D)** images (light grey: PART-Braak II; blue grey: AD-Braak IV; dark grey: AD-Braak stage VI). Red dashed lines indicate the area-cut off used for classifying tau aggregates: nano-aggregates (AT8 immunolabeled: 0.006 – 0.017 $\mu m^2$ ; AT270 immunolabeled: 0.004 – 0.017 $\mu m^2$), intermediate-aggregates (0.017 – 0.15 $\mu m^2$) and micro-aggregates (above 0.15 $\mu m^2$). Solid line indicates median, and the dotted lines indicate the 25th and 75th percentile. An unpaired, non-parametric Mann-Whitney test was performed between PART and AD-stages IV and VI. A p value of < 0.05 was taken as statically significant. . A p value of < 0.05 was taken as statically significant. P values = (ns) >0.05, (*) 0.05 – 0.03, (**) 0.002 – 0.03, (***) 0.0002 – 0.002, (****) 0.0001 – 0.0002.

**Fig. S3.**

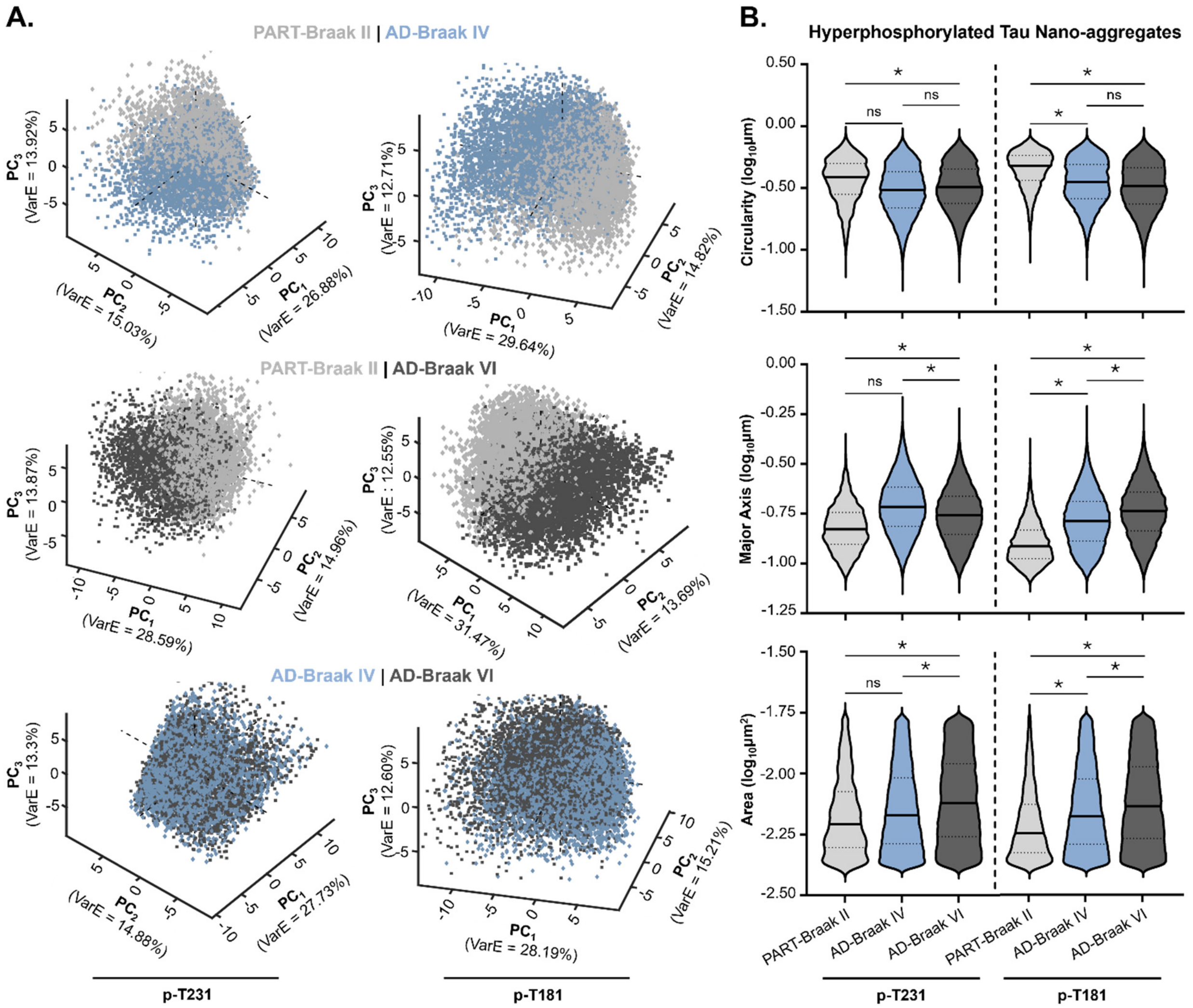


**Morphological analysis of p-T231 and p-T181 positive tau nano-aggregates. (A)** Principal Component Analysis (PCA) plots showing the PC 1 and 2 for p-T231 and p-T181 nano-aggregates in PART-Braak II (light grey dots), AD-Braak IV (blue grey dots) and AD-Braak VI (dark grey dots). **(B)** Violin plots showing the circularity, major axis, and area for p-T231 and p-T181 nano-aggregates in PART-Braak II (light grey dots), AD-Braak IV (blue grey dots) and AD-Braak VI (dark grey dots). Solid line indicates median, and the dotted lines indicate the 25th and 75th percentile. **p-T231:** PART-Braak II: N = 4 tissue sections, n = 15 fields of view; AD-Braak IV: N = 4 tissue sections, n = 18 fields of view; AD-Braak VI: N = 4 tissue sections, n = 17 fields of view. **p-T181** PART-Braak II: N = 4 tissue sections, n = 17 fields of view; AD-Braak IV: N = 4 tissue sections, n = 19 fields of view; AD-Braak VI: N = 4 tissue sections, n = 18 fields of view. A Wilcoxon rank sum test was performed between populations of tau aggregates in PART and AD-Braak IV and VI. A p value of < 0.05 was taken as statically significant. P values = (ns) >0.05, (*) 0.05 – 0.01, (**) < 0.01.

Fig. S4.

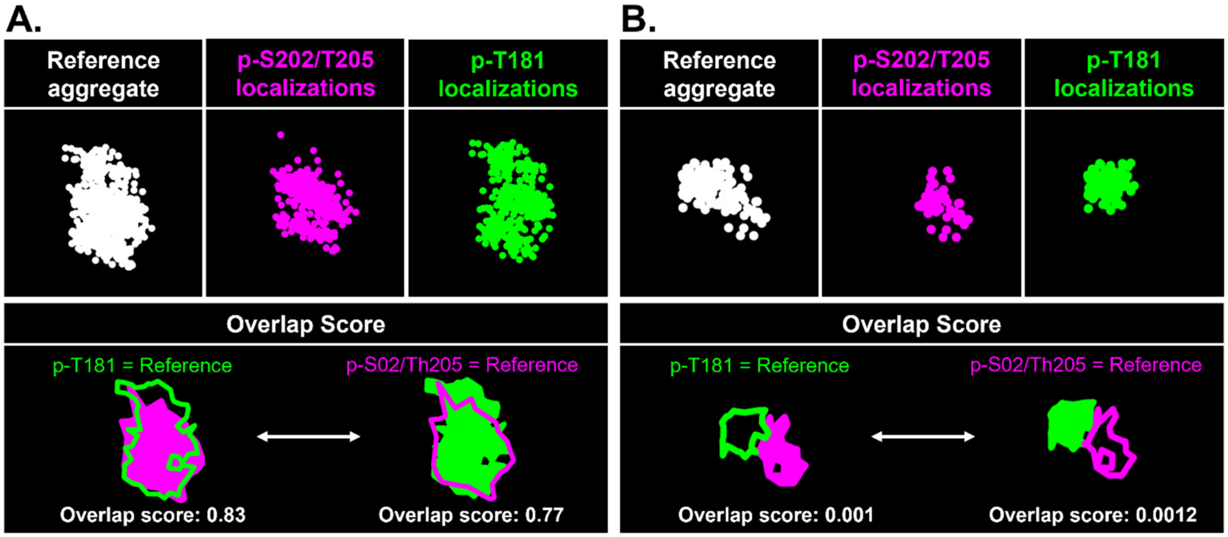


**Quality control step for segmentation in dual-color DNA-PAINT images.** Representative example of a properly segmented tau nano-aggregate with high overlap score **(A)** and a segmentation error with low overlap score **(B)**. White shows the reference tau aggregate localized points after segmentation from the reference merged image. Magenta shows the localized points corresponding to the p-S202/T205 channel. Green shows the localized points corresponding to the p-T181 channel. Magenta and green outlines show the alpha-shape computed from each channel overlaid on top of the other channel. A high overlap with the alpha-shape gives rise to a high overlap score and vice versa.

**Fig. S5.**

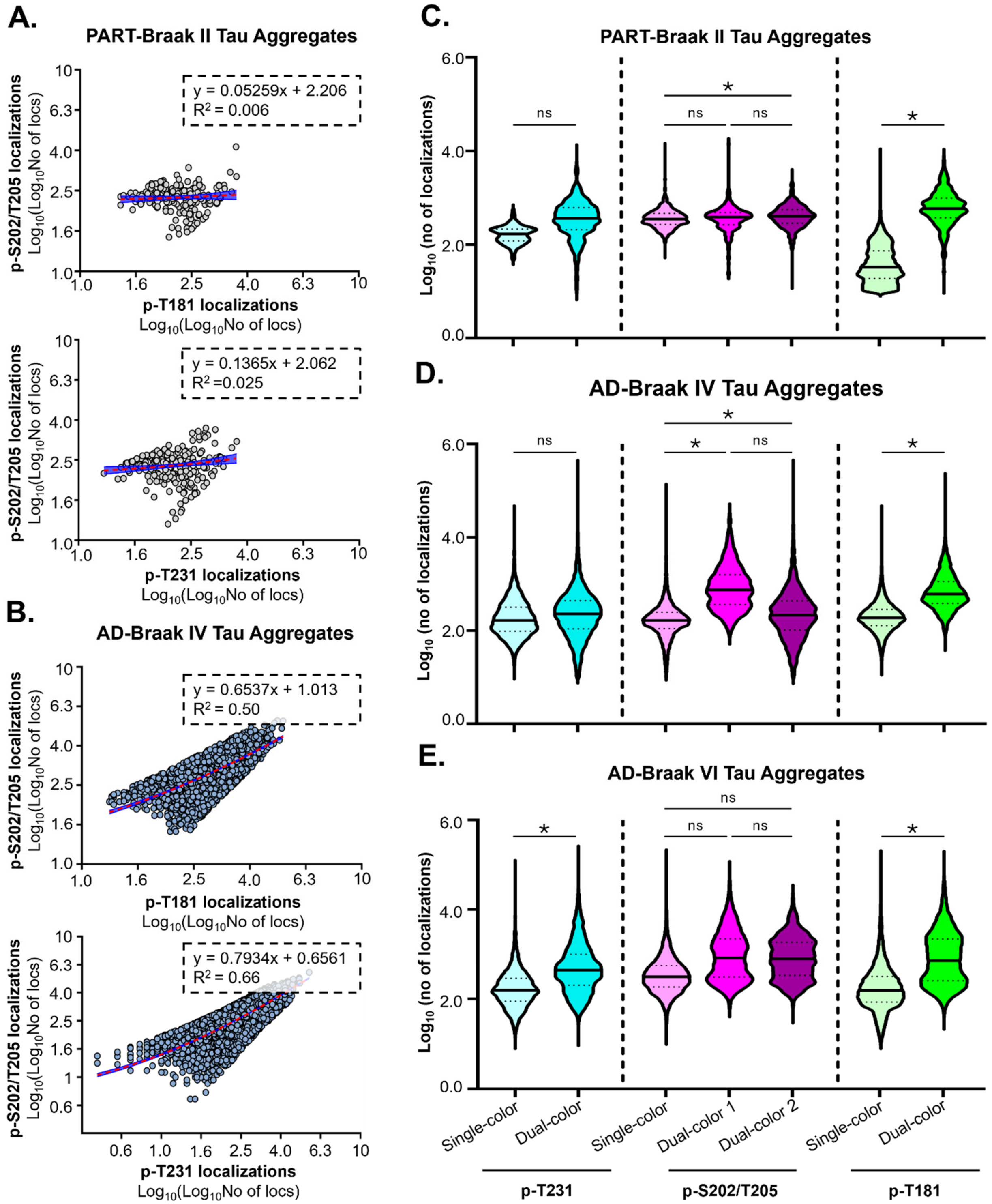


**Multicolor DNA-PAINT imaging controls. (A,B)** X,Y scatter plot showing the number of localizations of p-S202/T205 and p-T181 or p-S202/T205 and p-T231 per tau aggregates in PART-Braak II **(A)** and AD-Braak IV **(B)** tissue sections. Red line is the linear regression, blue shadow is the standard error. AT8 + AT270 immunolabeled PART-Braak II and AD-Braak IV tissue sections: N = 4 tissue sections, n = 16 fields of view; AT8 + AT180 immunolabeled PART-Braak II and AD-Braak IV tissue sections: N = 4 tissue sections, n = 16 fields of view. **(C-E)** Violin plots show the number of localizations per tau aggregate for p-T231 (cyan), p-S202/T205 (magenta) and p-T181 (green) computed from single-color DNA-PAINT images (light color) or dual-color DNA-PAINT images (dark color) in PART-Braak II **(C)**, AD-Braak IV **(D)** and AD-Braak VI **(E)**. p-S202/T205 has two plots for dual-color images corresponding to the two combinations for p-S202/T205 with p-T181 (magenta) and pS202/T205 with p-T231 (purple), respectively. Single-color DNA-PAINT images: **p-T231:** PART-Braak II: N = 4 tissue sections, n = 15 fields of view; AD-Braak IV: N = 4 tissue sections, n = 18 fields of view; AD-Braak VI: N = 4 tissue sections , n = 17 fields of view. **p-S202/T205:** PART-Braak II: N = 4 tissue sections, n = 17 fields of view; AD-Braak IV: N = 4 tissue sections, n = 17 fields of view; AD-Braak VI: N = 4 tissue sections, n = 18 fields of view. **p-T181:** PART-Braak II: N = 4 tissue sections, n = 17 fields of view; AD-Braak IV: N = 4 tissue sections, n = 19 fields of view; AD-Braak VI: N = 4 tissue sections, n = 18 fields of view. Dual-color DNA-PAINT images: **p-T231**: PART-Braak II: N = 4 tissue sections, n = 16 fields of view; AD-Braak IV: N = 4 tissue sections, n = 16 fields of view; AD-Braak VI: N = 4 tissue sections, n = 16 fields of view.; **p-S202/Thr205**: PART-Braak II: N = 8 tissue sections, n = 32 fields of view; AD-Braak IV: N = 4 tissue sections, n = 32 fields of view; AD-Braak VI: N = 8 tissue sections, n = 32 fields of view; **p-T181**: PART-Braak II: N = 4 tissue sections, n = 16 fields of view; AD-Braak IV: N = 4 tissue sections, n = 16 fields of view; AD-Braak VI: N = 4 tissue sections, n = 16 fields of view. An unpaired, non-parametric Mann-Whitney test was performed between PART and AD-stages IV and VI. A p value of < 0.05 was taken as statically significant. P values = (ns) >0.05, (*) 0.05 – 0.03, (**) 0.002 – 0.03, (***) 0.0002 – 0.002, (****) 0.0001 – 0.0002.

**Fig. S6**

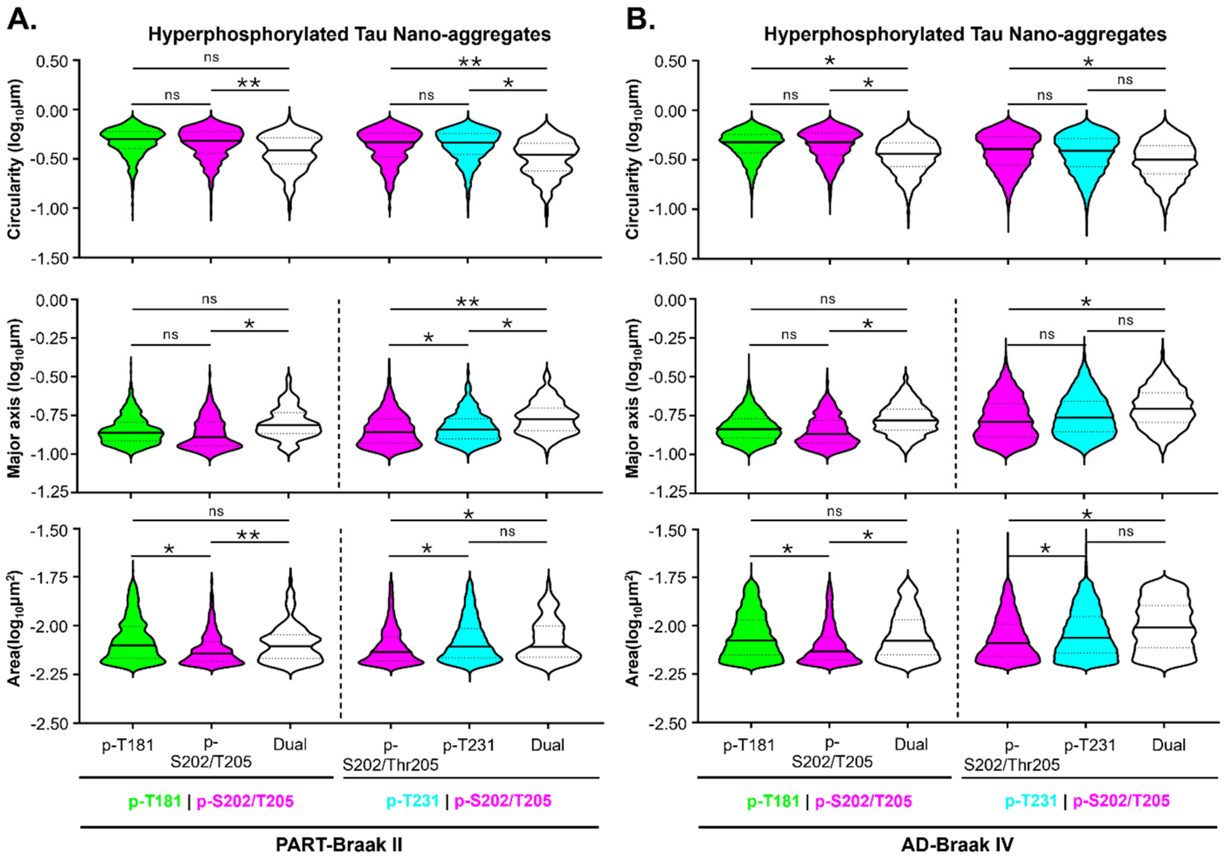


**Morphological features of tau nano-aggregates from PART-Braak II and AD-Braak IV. (A, B)** Violin plots showing the circularity, major axis, and area for p-T181 only (green), p-S202/T205 only (magenta), p-T231 only (cyan), or p-T181/p-S202/T205 and p-T231/p-S202/T205 dually modified (white) hyperphosphorylated tau nano-aggregates present in dual-color DNA-PAINT images from PART-Braak II **(A)** or AD-Braak IV **(B)**. Solid line indicates median, and the dotted lines indicate the 25th and 75th percentile. **AT8 + AT270** immunolabeled tissue sections: N = 4 tissue sections, n = 16 fields of view for both PART-Braak II and AD-Braak IV; **AT8 + AT180** immunolabeled tissue sections: N = 4 tissue sections, n = 16 fields of view for both PART-Braak II and AD-Braak IV. A Wilcoxon rank sum test was performed between enrichment score specific populations of tau aggregates in AD-stage VI. A p value of < 0.05 was taken as statically significant. P values = (ns) >0.05, (*) 0.05 – 0.01, (**) < 0.01.
